# Depletion of lamin-associated polypeptide 2alpha leads to chromatin reorganization and binding of A-type lamins to open genomic regions

**DOI:** 10.64898/2026.08.03.742457

**Authors:** Daria Filipczak, Fatih Sarigol, Daniel Malzl, Roland Foisner, Nana Naetar

**Author notes:** **Corresponding authors:** Nana Naetar,**, Corresponding author, Lead contact:** Roland Foisner. These authors contributed equally to this work (Daria Filipczak, Fatih Sarigol).

## Abstract

**Background:** Lamins are major regulators of the spatial and functional organization of chromatin. Lamins at the nuclear periphery form the lamina that anchors heterochromatin to the nuclear envelope. A subpool of A-type lamins localizes in the nuclear interior, where they also bind to euchromatic genomic regions. A-type lamin properties and chromatin association are regulated by lamin-associated polypeptide 2alpha (LAP2α). Here we systematically analyze, how LAP2α depletion affects chromatin organization, accessibility and gene expression on a genome-wide level.

**Results:** LAP2α depletion in mouse dermal fibroblasts positively and negatively affects chromatin accessibility and gene expression throughout the genome, which correlates with changes in chromatin association of A-type lamins and the nucleosomal remodeler proteins BRG1 and CHD4. In particular, A-type lamins bind to open chromatin regions close to BRG1 and CHD4 binding sites and deregulated genes, but do not directly accumulate on genes and BRG1 and CHD4-enriched sites. Unsupervised clustering of the datasets on LAP2α-bound genomic regions confirms spreading of A-type lamins to active chromatin regions containing deregulated genes and an enrichment of chromatin remodelers on a subset of these genomic regions.

**Conclusions:** LAP2α depletion in fibroblasts leads to a gross rearrangement of chromatin. Genome-wide chromatin reorganization is linked to spreading of A-type lamins to active chromatin regions and accompanied by a restriction of chromatin remodelers to a subset of active genomic regions. These changes correlate with changes in chromatin accessibility and gene expression throughout the genome, particularly in regions where lamin binding is gained in LAP2α knockout versus wildtype cells.

## Background

The spatial organization of chromatin within the nucleus is fundamental to gene regulation and is characterized by chromatin compartmentalization, with heterochromatin predominantly localized at the nuclear periphery and euchromatin distributed throughout the nuclear interior [1].

Lamins are major regulators of chromatin organization and function. At the nuclear periphery, lamins form a filamentous protein meshwork underneath the inner nuclear membrane, called the nuclear lamina [2]. The lamina provides mechanical support to the nucleus, fulfills essential functions in mechanosensing and mechanosignaling [3–5] and has an essential role in the organization and regulation of chromatin. It serves as an anchor for heterochromatin through association with lamina-associated domains (LADs), large transcriptionally inactive genomic regions of megabase scale enriched in repressive histone marks [6]. In mammalian cells, the nuclear lamina comprises four main lamin isoforms: lamin A, lamin C, lamin B1 and lamin B2 and numerous lamin-and chromatin-interacting proteins of the inner nuclear membrane [7]. While B-type lamins B1 and B2 are tightly attached to the inner nuclear membrane and predominantly located at the nuclear periphery, A-type lamins (A and C), encoded by the *LMNA* gene, exist in two distinct nuclear subpools: stable filaments as part of the lamina at the nuclear periphery interacting mostly with heterochromatic LADs, and a dynamic pool in the nuclear interior that associates also with euchromatin [8].

Lamin-associated polypeptide 2 alpha (LAP2α), a nucleoplasmic protein encoded by one of six splice variants of the *TMPO* gene [9] is a key regulator of the cellular localization, structure and functions of A-type lamins. Like all other splice variants, LAP2α contains a LAP2-Emerin-Man1 (LEM) domain, a 40 residues long helix-loop-helix domain interacting with the small DNA-binding protein barrier-to-autointegration factor (BAF) [10, 11]. The LEM domain mediates the interaction of LAP2α with chromatin in cooperation with the lamin A/C-interaction domain and several additional regions in the polypeptide [12]. LAP2α depletion has been shown to significantly impact A-type lamin dynamics. Specifically, loss of LAP2α increases lamin-chromatin association and promotes the formation of higher-order lamin structures that exhibit reduced mobility and increased resistance to biochemical extraction [8, 12]. LAP2α regulates lamin A/C interactions with chromatin by competing for chromatin binding sites and by forming different complexes with lamins and BAF [12]. These complexes can associate with both euchromatic and heterochromatic regions and likely affect chromatin organization and gene regulation [8, 13]. Moreover, LAP2α has been shown to facilitate myogenic gene expression by preventing lamin spreading to active chromatin regions containing a subset of myogenic genes [14]. Probably through these lamin A/C-regulating functions, LAP2α may also affect the transition of cells from a quiescent to a proliferative or differentiation state as previously shown in various cell systems [15, 16] and in mice [17]. Studies in LAP2α knockout mice have demonstrated that LAP2α deficiency delays the differentiation of progenitor cells in muscle and other regenerative tissues and affects tissue homeostasis [15, 16, 18, 19].

While we have previously found that LAP2α affects the assembly state and mobility of nucleoplasmic A-type lamins [8] and decreases lamin chromatin association [12] and influences gene expression during musle differention [14], this study aimed to investigate comprehensively the broader consequences of LAP2α depletion on genome-wide chromatin organization, accessibility and gene expression in the context of chromatin association of nucleosomal remodelers, which are involved in the regulation of chromatin accessibility and gene expression. Specifically, we examined the chromatin remodelers BRG1 and CHD4, which have been shown to associate with lamins in various nuclear contexts [20]. BRG1, a subunit of the SWI/SNF remodeling complex, facilitates gene expression through nucleosome repositioning, while CHD4, a component of the NuRD complex, serves dual functions as both a nucleosomal remodeler on active chromatin and a histone deacetylase, primarily targeting transcriptionally silent chromatin regions [21].

Using a previously generated and characterized isogenic pair of immortalized LAP2α wildtype and LAP2α knockout mouse dermal fibroblasts [8], we performed genome-wide analyses including ChIP-seq, RNA-seq and ATAC-seq as well as unsupervised genome clustering, to systematically study changes in chromatin accessibility, gene expression and chromatin organization following LAP2α depletion. We correlated these changes with alterations in the chromatin association of lamin A/C and of chromatin remodelers. Our results reveal that LAP2α-enriched genomic regions undergo substantial rearrangements upon LAP2α depletion, leading to the binding of lamins to active chromatin regions, which may in turn restrict the localization of chromatin remodelers to a subset of active loci. These rearrangements are accompanied by changes in chromatin accessibility and gene expression particularly in lamin A/C-bound regions. Altogether, this study provides new insights into the regulatory functions of LAP2α on a genome-wide level, highlighting its role in restricting lamin A/C binding to open chromatin and thereby potentially enabling access of chromatin regulators.

## Results

### LAP2α loss induces genome-wide changes in gene expression and chromatin accessibility

We have previously shown that depletion of LAP2α in myoblasts and fibroblasts leads to changes in chromatin association of lamin A/C [12–14]. Here we aimed at elucidating the effects of altered lamin A/C chromatin association upon LAP2α loss on chromatin organization and accessibility and gene expression on a genome-wide level. We used immortalized mouse dermal fibroblasts and an isogenic LAP2α knockout fibroblast cell line generated previously by CRISPR/Cas9 mediated knockout of LAP2α [8]. First, we assessed genome-wide changes in chromatin accessibility in LAP2α knockout versus wildtype cells by ATAC-seq combined with differential accessibility analysis. We identified 10,139 genomic regions with significantly decreased accessibility and 6,828 regions with significantly increased accessibility in cells lacking LAP2α compared to wildtype cells (Fig. 1A). These differentially accessible regions had an average length of 403 bp and were distributed across all chromosomes (Fig. 1B). It remains unclear why chromosomes 18 and 19 seem to have a slightly different enrichment of accessible regions compared to the other chromsomes, as they do not differ significantly from other chromosomes in gene density (GRCm38.102 from ENSEMBL) and amounts of repeatitive sequences (UCSC browser; Group: Variation and Repeats. Track: RepeatMasker), and LADs and interLAD regions (GEO, ID GSE17051).

**Figure 1.**
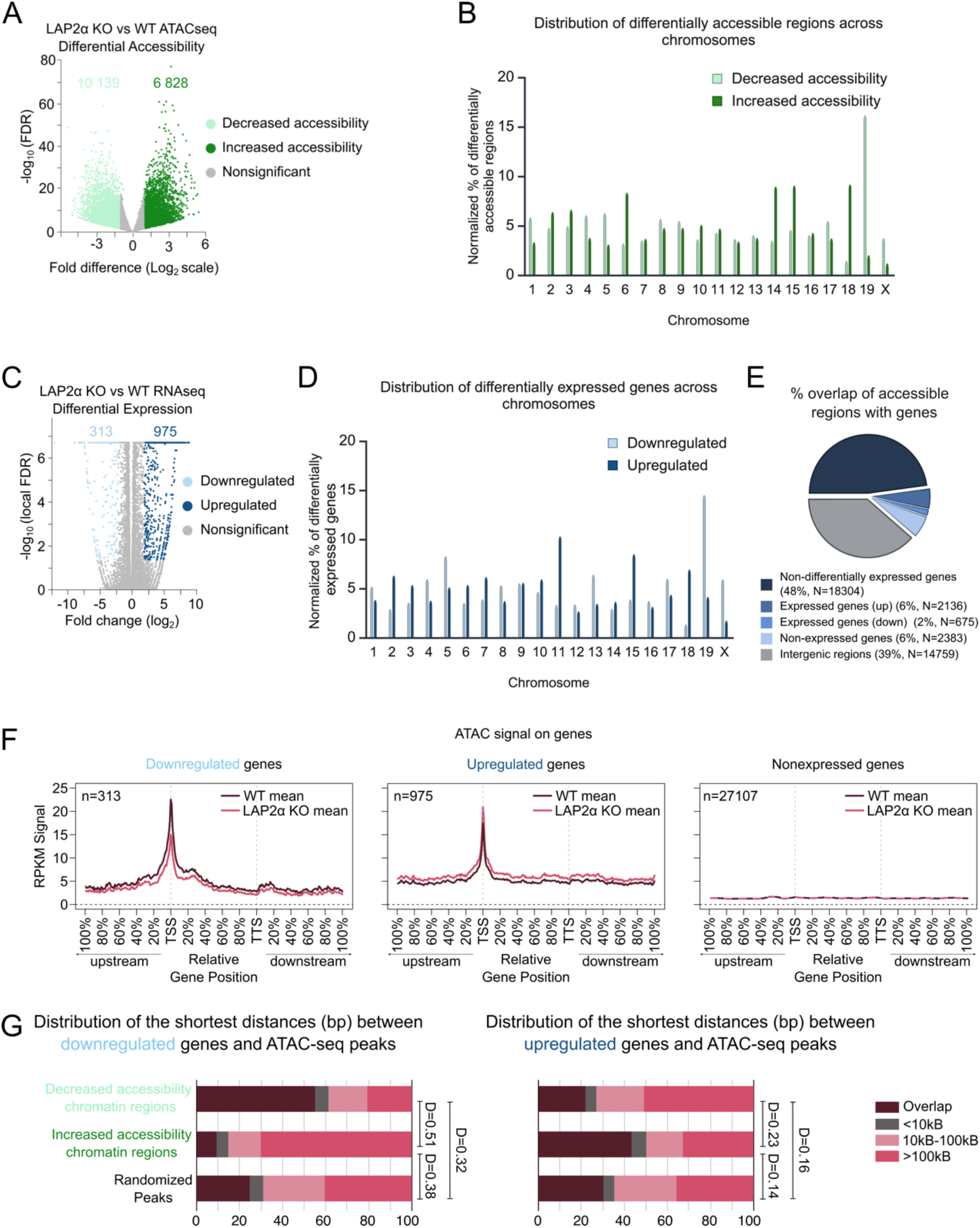
Depletion of LAP2α causes changes in chromatin accessibility and gene expression. **(A)** ATAC-seq analysis in LAP2α knockout versus wildtype fibroblasts. The volcano plot shows differentially accessible regions in LAP2α knockout versus wildtype fibroblasts (minimum difference of 1 in log2-transformed values of accessibility, FDR < 0.01). Regions with decreased accessibility are shown in light green, regions with increased accessibility are shown in dark green and regions with no significant change in accessibility are shown in gray. **(B)** Distribution of differentially accessible regions across chromosomes. Bar chart displays the percentage of regions localized on each chromosome, normalized to chromosome length. **(C)** Analysis of gene expression changes in LAP2α knockout versus wildtype fibroblasts. The volcano plot displays differentially expressed genes in LAP2α knockout versus wildtype fibroblasts (minimum log2 fold change of 2, local FDR<0.05, baseMean>5). Downregulated genes are shown in light blue, upregulated genes in dark blue and genes with no significant changes in expression are shown in gray. **(D)** Distribution of differentially expressed genes across chromosomes. Bar chart displays the percentages of genes localized on each chromosome, normalized to chromosome lengths. **(E)** Pie chart showing the percentage overlap of all accessible chromatin regions with genes and intergenic regions. **(F)** Mean RPKM ATAC-seq signal from wildtype (WT; brown) and LAP2α knockout (KO; pink) cells plotted on downregulated (left panel), upregulated (middle panel) and non-expressed (right panel) genes. **(G)** Bar charts showing genomic distances in base pairs between differentially accessible regions (split into regions with decreased and increased accessibility) and the closest downregulated (left panel) or upregulated (right panel) genes. Distances of randomized ATAC-seq peaks to genes are included as a control. Differences between distributions are indicated by the D-value from the KS test, representing the maximum distance between the cumulative distributions of the two sets. All KS tests have an adjusted P-value (P adj) below 0.05.

To see, if and how these changes in chromatin accessibility in LAP2α knockout versus wildtype cells correlate with changes in gene expression we performed RNA-seq. This analysis revealed 313 significantly downregulated and 975 significantly upregulated genes in LAP2α knockout versus wildtype cells throughout the genome (Fig. 1C, D). 8% of the accessible genomic regions in LAP2α knockout and wildtype cells overlapped with differentially expressed genes in LAP2α knockout versus wildtype cells (Fig. 1E). The majority of accessible sites, however overlapped with expressed genes (48%) while only 6% overlapped with non-expressed genes (Fig. 1E). Interestingly, a large fraction of accessible sites (39%) overlapped also with intergenic regions. As expected, the ATAC-seq signal was slightly reduced on the transcription start site (TSS) of downregulated genes in LAP2α knockout versus wildtype cells and was un-changed or slightly increased on the TSS of upregulated genes, while non-expressed genes were not accessible (Fig. 1F). Furthermore, by testing the closest distances between differentially expressed genes and differentially accessible ATAC-seq peaks we found that around 60% of regions with significantly reduced accessibility overlapped with downregulated genes or were closer than 10kb, while only around 15% of the regions with increased accessibility and 30% of randomized genomic regions located within 10 kb of the downregulated genes (Fig. 1G, left panel). Conversely, about 50% of genomic regions showing significantly increased accessibility overlapped (+/-10kB) with upregulated genes, compared to only 20% of regions with decreased accessibility and 30% of randomized genomic regions (Fig. 1G, right panel). Thus, genomic regions with reduced accessibility are enriched in downregulated genes, those with increased accessibility in upregulated genes, clearly above the number of randomly distributed genes.

Overall, these results show that depletion of LAP2α leads to gross changes in both gene expression and chromatin accessibility and that these changes correlate closely with each other. However, besides accessibility changes on or around genes, many intergenic genomic regions show changes in accessibility, pointing towards a global reorganization of chromatin structure upon LAP2α depletion.

### Chromatin association of lamin A/C and nucleosomal remodeler proteins changes upon depletion of LAP2α

Having observed that the loss of LAP2α causes genome-wide changes in chromatin accessibility and gene expression, we next investigated how these changes relate to the redistribution of lamin A/C on chromatin. In addition, we tested genome-wide chromatin association of components of the SWI/SNF and NuRD chromatin remodelers because these complexes are known to regulate chromatin accessibility by moving and/or ejecting nucleosomes on gene regulatory regions [22]. In particular, we tested chromatin association of BRG1, an ATPase of the SWI/SNF complex and CHD4, a component of NuRD, as these proteins were proposed to bind LAP2α in proximity-based proteome-wide interaction analyses (see [23]). Using co-immunoprecipitation assays, we confirmed binding of BRG1 and CHD4 to endogenous full length LAP2α and to ectopic full length LAP2α expressed in LAP2α knockout cells (Additional file 1: Fig. S1A, B). As a negative control, N-and C-terminal truncation mutants of LAP2α did not interact with the remodelers confirming specificity of the interactions. (Additional file 1: Fig. S1A, B, C). Because the N-and C-terminal domains of LAP2α were previously found to jointly mediate the interaction of LAP2α with chromatin and lamin A/C [12], the lack of binding of BRG1 and CHD4 to LAP2α truncation mutants suggests that both domains are also involved in the interaction with nucleosomal remodelers. Additionally, BRG1 and CHD4 coprecipitate with lamin C and to a lesser extent with lamin A. (Additional file 1: Fig. S1A, B). However, association of LAP2α and lamin A/C with nucleosomal remodeler complexes seems to be independent of each other, because knockout of either protein does not affect binding of the other (Additional file 1: Fig. S1A, B).

Most previous studies analyzing chromatin association of lamin A/C and LAP2α on a genome-wide level used the enhanced domain detector (EDD) peak caller [24], which detects lamin enrichment over large Mb scale genomic regions (Additional file 2: Table S1) in both heterochromatic LADs and in inter-LAD regions (Fig. 2A, EDD peaks). However, as BRG1 and CHD4 are known to bind to smaller genomic regions (<500 bp) [25, 26], we aimed at testing lamin A/C chromatin association at a smaller scale (300-500 bp) using a combination of the MACS2 and SPP peak calling algorithms with parameterization (see methods for details) (Additional file 2: Table S1). This approach allowed us to directly compare the genome-wide chromatin binding of lamin A/C and LAP2α with that of BRG1 and CHD4. Furthermore, in order to improve ChIP efficiency and enhance detection of BRG1 and CHD4, we performed dual cross-linking ChIP (protein-protein followed by protein-DNA crosslinking using disuccinimidyl glutarate (DSG) and formaldehyde, respectively) and applied two sonication conditions (16 and 20 sonication cycles of sonication) to include both heterochromatic genomic regions, enriched in LADs and non-expressed genes (Additional file 1: Fig. S2A, B, 20 sonication cycles), and euchromatic regions, enriched in ciLADs and expressed genes (Additional file 1: Fig. S2A, B, 16 sonication cycles). Although the IGV browser tracks of ChIP signals in the 16 and 20 sonication cycles samples look similar, the overlap of MACS2/SPP peaks obtained from the 16 and 20 sonication cycles ChIP samples was low, particularly for LAP2α and lamin A/C, confirming the enrichment of different genomic regions in these samples (Additional file 1: Fig. S2C, D). In order to test the feasibility of the double cross-linking ChIP also for analyzing lamin A/C chromatin association, which was predominantly analyzed using the classical single crosslinking ChIP protocol, we compared previously published results obtained in single formaldehyde cross-linking lamin A/C ChIP analyses [13] with data obtained in our ChIP protocol using double crosslinking. In agreement with the published data, we found significant overlap of lamin A/C-and LAP2α-enriched genomic regions detected by the EDD peak caller, as well as a redistribution of lamin A/C on chromatin in LAP2α knockout cells (see three-way Venn diagram comparing lamin A/C EDD peaks in wildtype and LAP2α knockout cells and LAP2α EDD peaks in wildtype cells, Additional file 1: Fig. S2E). Thus, dual-crosslinking ChIP provides results consistent with previous studies for lamins and LAP2α. In addition, we tested reproducibility of the lamin A/C ChIP by using two different lamin A/C antibodies (3A6 and E1). The Pearson correlation of lamin A/C ChIP signals for these two antibodies is 0.91 for the 16 sonication cycles samples and 0.94 for the 20 cycles samples, confirming high reproducibility of lamin A/C ChIP with the double cross linking ChIP protocol.

**Figure 2.**
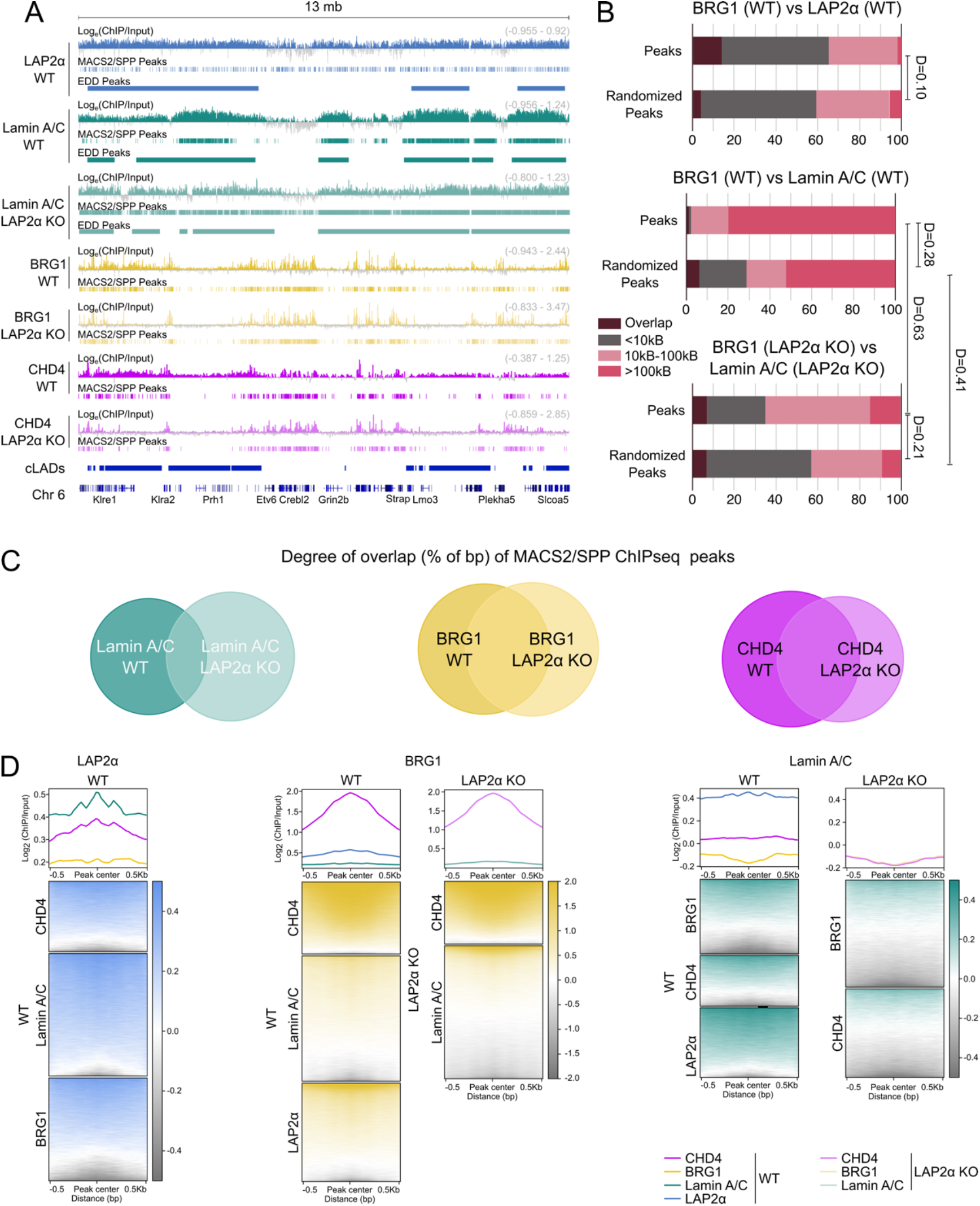
Depletion of LAP2α causes redistribution of lamin A/C and chromatin remodelers on chromatin. **(A)** ChIP-seq analysis was performed in wildtype and LAP2α knockout fibroblasts for LAP2α (blue), lamin A/C (3A6 antibody; turquoise), BRG1 (yellow) and CHD4 (magenta) as indicated. The IGV browser was used to display the log2 ratio of ChIP over input signal from samples sonicated for 16 sonication cycles, with tracks shown for mouse chromosome 6. Positive log2 ratio values are shown in color, while negative values are shown in gray. Peaks called by the MACS2/SPP peak callers and corrected with HOMER and by the EDD peak caller (for lamin A/C and LAP2α) for samples sonicated for 16 and 20 sonication cycles were merged and are depicted for each ChIP track. The scale of each log2 ratio track is indicated on the right. Constitutive lamina-associated domains (cLADs) are also annotated. Gene annotations are based on the NCBI reference sequence database. **(B)** Bar charts displaying genomic distances in base pairs between ChIP-seq peaks for LAP2α and BRG1 (upper panel), lamin A/C and BRG1 (middle panel) in wildtype cells and lamin A/C and BRG1 (lower panel) in LAP2α knockout cells. Randomized peaks are included as controls. LAP2α peaks were randomized in the upper panel, lamin A/C peaks in lower and middle panels. Differences between distributions are indicated by the D-value from the KS test, representing the maximum distance between the cumulative distributions of the two sets. All KS tests have an adjusted P-value (P adj) below 0.05. **(C)** Venn diagrams showing the percentage of basepair overlap (% of bp) for ChIP-seq peaks of proteins in wildtype and LAP2α knockout cells: lamin A/C (left diagram), BRG1 (middle diagram) and CHD4 (right diagram). **(D)** Heatmaps showing log2 ratio signals (ChIP over input) on combined (16 and 20 sonication cycles) ChIP-seq peaks +/-0.5kb from peak center. Log2 ratio signals (16 sonication cycles) are presented as follows: LAP2α in wildtype fibroblasts on peaks of BRG1, lamin A/C and CHD4 (left panel); BRG1 in wildtype and LAP2α knockout fibroblasts on peaks of CHD4, lamin A/C and LAP2α (middle panel); lamin A/C in wildtype and LAP2α knockout fibroblasts on peaks of BRG1, CHD4 and LAP2α (right panel). Graphs above the heatmaps display the mean log2 ratio signals.

Using this double cross-linking ChIP protocol in combination with the MACS2/SPP peak caller, we identified enrichment of LAP2α and lamin A/C in many individual short MACS2/SPP peaks often located within large EDD peaks and overlapping with heterochromatic LADs. In addition, lamin A/C and particularly LAP2α also showed a number of MACS2/SPP peaks outside of heterochromatic LADs in genomic regions not detected by the EDD peak caller (Fig. 2A). In contrast, BRG1 and CHD4 were predominantly enriched in short, well-defined regions, mostly located outside of heterochromatic LADs, but also to some degree within LADs (Fig. 2A).

In order to identify and analyse the enrichment of the proteins on total chromatin, we combined peaks obtained in the 16 and 20 sonication cycles ChIP samples, which are enriched in different chromatin fractions (Additional file 1: Fig. S2C, D). Closest distance analyses revealed that about 60% of BRG1 and CHD4 peaks were located within 10 kb or overlapped with LAP2α-bound regions, similar to randomized peaks (Fig. 2B, Additional file 1: Fig. S3A). In contrast, the majority of lamin A/C peaks were more than 100 kb away from BRG1 or CHD4 peaks in wildtype cells, a distance significantly larger than that of randomized peaks. This suggests that unlike LAP2α lamin A/C largely avoids genomic regions occupied by BRG1 or CHD4 (Fig. 2B, Additional file 1: Fig. S3A).

Interestingly, we observed a redistribution of lamin A/C, BRG1 and CHD4 on chromatin following LAP2α depletion, with lamin A/C showing the most pronounced change, as revealed by analyzing the overlap of peaks in LAP2α knockout versus wildtype cells (Fig. 2C). Additionally, while the majority of BRG1 and CHD4 peaks were more than 100 kb away from lamin A/C peaks in wildtype cells, 35% of BRG1 and 70% of CHD4 peaks were within 10 kb or overlapped with lamin A/C peaks in LAP2α knockout cells, similar to the distance to randomized peaks, suggesting that lamin A/C, BRG1 and CHD4 do no longer avoid each other but move closer on chromatin in LAP2α knockout compared to wildtype cells (Fig. 2B, Additional file 1: Fig.S3A). In contrast, 80% of BRG1 and CHD4 peaks overlapped or were located within 10kb distance in wildtype cells and this was unchanged in LAP2α knockout cells (Additional file 1: Fig. S3A), suggesting that BRG1 and CHD4 locate to similar genomic regions. Thus, loss of LAP2α leads to gross changes in the binding of lamin A/C to chromatin bringing lamins into close proximity with chromatin remodelers.

Although lamin A/C moved closer to BRG1-and CHD4-bound genomic regions in LAP2α knockout versus wildtype cells, the lamin A/C ChIP signal was not significantly enriched on BRG1 or CHD4 bound regions in wildtype and LAP2α knockout cells in both the 16 and 20 sonication cycles samples, while the BRG1 and the CHD4 signals coincided (Fig. 2D, Additional file 1: Fig.S3B, C). In contrast to lamin A/C, the LAP2α signal was slightly increased on BRG1 and CHD4 peaks, but did not reach the level of enrichment seen on lamin A/C peaks in wildtype cells (Fig. 2D, Additional file 1: Fig.S3B, C).

In summary, these results show that, unlike lamin A/C-bound regions, genomic regions bound by LAP2α are in close proximity or overlap with regions occupied by BRG1 and CHD4 in wildtype cells. When LAP2α is depleted, lamin A/C mostly redistributes on chromatin, moving closer to BRG1 and CHD4, but does not directly enrich on BRG1-and CHD4-bound genomic regions and vice versa.

### Redistribution of lamin A/C on chromatin correlates with gene expression changes and to a lesser extent with accessibility changes

To investigate, if and how the redistribution of lamin A/C, BRG1 and CHD4 on chromatin relates to changes in chromatin accessibility and gene expression in LAP2α knockout versus wildtype cells, we first examined the overlap of ChIP-seq peaks for LAP2α, Lamin A/C, BRG1 and CHD4 with the genomic regions showing significantly increased or decreased accessibility.

BRG1 and CHD4-bound regions overlapped with a large fraction (60–80%) of differentially accessible sites in wild-type cells clearly above the overlap with randomized sites. The overlap of BRG1 and CHD4 peaks with these sites was reduced in regions of decreased accessibility and increased at regions of higher accessibility in LAP2α knockout versus wildtype cells (Fig. 3A). In contrast, LAP2α and lamin A/C showed less than 20% overlap with differentially accessible sites in wildtype cells, which is similar to randomized sites, confirming that lamin A/C binds mostly to non-accessible regions in wildtype cells. The overlap of lamin A/C-bound regions with differentially accessible regions, however increased two-to three-fold in LAP2α knockout cells, supporting the hypothesis that lamin A/C spreads to regions closer to accessible sites upon LAP2α knockout (Fig. 3A). Interestingly, closest distance analyses revealed that 80% of differentially accessible genomic regions were located within 10kb distance to regions that gained lamin A/C binding in LAP2α knockout cells. In contrast, the distance between differentially accessible regions and genomic sites that lost lamin A/C binding in LAP2α knockout versus wildtype cells was mostly larger than 10kb (Fig. 3B). We concluded that lamin A/C generally spreads to accessible genomic regions in LAP2α knockout versus wildtype cells. Heatmap analyses of the lamin A/C ChIP signals on differentially accessible regions revealed no enrichment of lamin A/C on differentially accessible regions in both 16 and 20 sonication cycles ChIP samples (Fig. 3C, Additional file 1: Fig.S4A). In sharp contrast, BRG1 and CHD4 showed an enrichment at differentially accessible regions, with a clearly increased BRG1 and CHD4 signal on regions with increased accessibility and a slightly reduced signal on regions with reduced accessibility in LAP2α knockout versus wildtype cells (Fig. 3C, Additional file 1: Fig.S4A). Thus, lamin A/C binds to genomic rgions in the vicinity of accessible sites in LAP2α knockout cells, but is not directly enrich on these sites.

**Figure 3.**
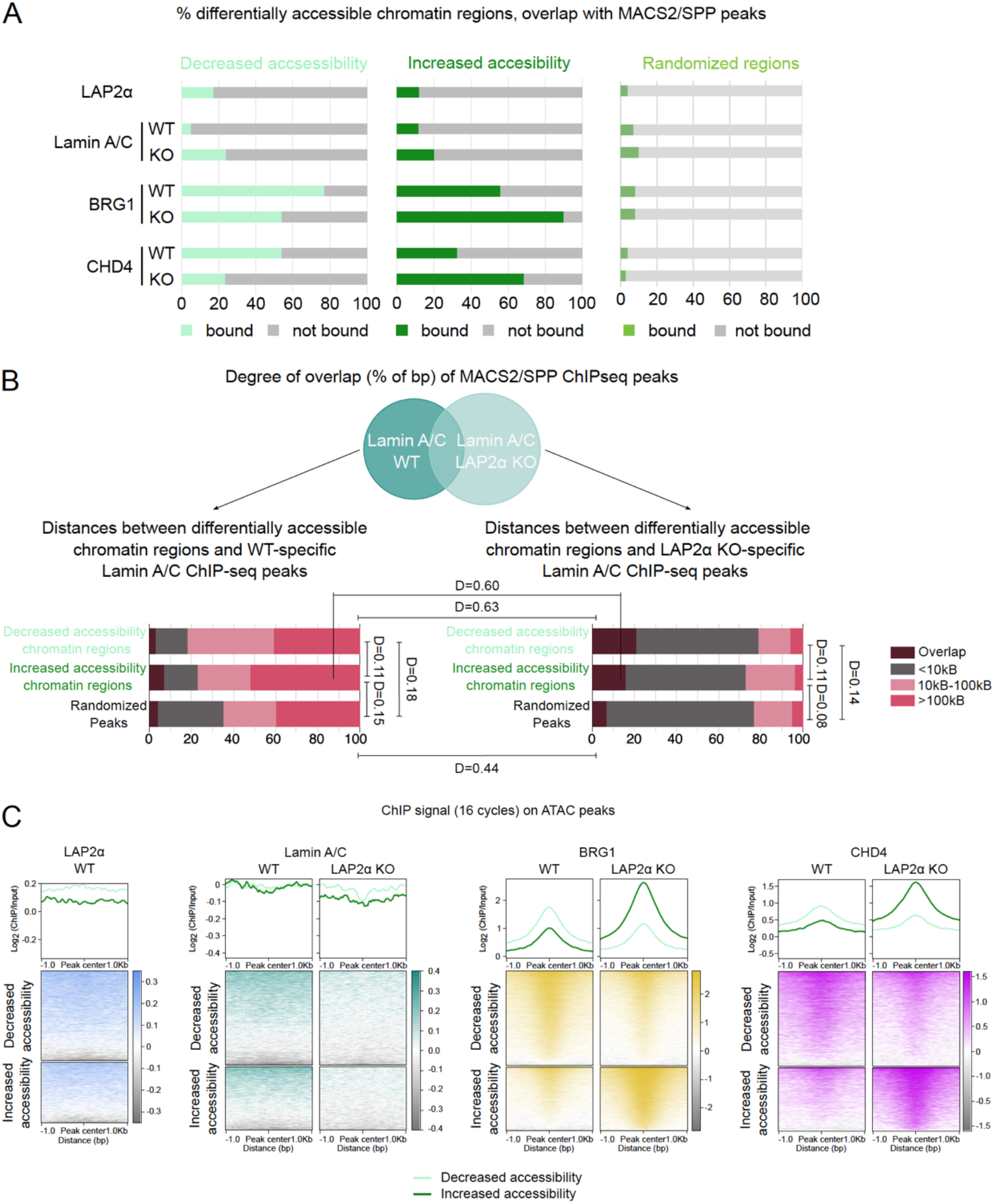
Changes in chromatin accessibility correlate with changes in chromatin association of chromatin remodelers. **(A)** Bar graphs showing the percentages of differentially accessible regions (decreased – left panel, increased – right panel) and 8480 randomized regions overlapping with combined (16 and 20 sonication cycles) ChIP-seq peaks of LAP2α in wildtype cells and lamin A/C, BRG1 and CHD4 in both LAP2α wildtype and knockout cells. **(B)** Venn diagrams showing the percentage of basepair overlap (% of bp) for ChIP-seq peaks of lamin A/C in wildtype and LAP2α knockout cells (upper panel). Bar charts (lower panel) showing genomic distances in base pairs between differentially accessible regions (split into regions with decreased and increased accessibility) and the closest lamin A/C ChIP-seq peaks, subdivided into peaks that are lost upon LAP2α depletion (left panel) and those that are gained (right panel). Distances of randomized ATAC-seq peaks to ChIP-seq peaks are included as a control. Differences between distributions are indicated by the D-value from the KS test, representing the maximum distance between the cumulative distributions of the two sets. All KS tests have an adjusted P-value (P adj) below 0.05. **(C)** Heatmaps showing log2 ratio signals (ChIP over input, 16 sonication cycles) on chromatin regions with decreased accessibility (light green) and increased accessibility (dark green) +/-1.0kb from center in LAP2α knockout versus wildtype cells. Log2 ratio signals are presented in the following order from left to right: LAP2α, lamin A/C, BRG1 and CHD4. Graphs above the heatmaps display the mean log2 ratio signals.

Next, we examined a potential correlation between the redistribution of lamin A/C, BRG1 and CHD4 on chromatin and the differential expression of genes in LAP2α knockout versus wildtype cells. 80% of both downregulated and upregulated genes overlapped with BRG1-and CHD4-bound genomic regions in wildtype cells and this was mostly unchanged in LAP2α knockout cells (Fig. 4A). This high overlap of BRG1 and CHD4 with differentially expressed genes was significantly higher than that of non-expressed genes confirming the enrichment of nucleosomal remodelers on expressed genes. LAP2α also had a larger overlap (around 60%) with these gene sets compared to non-expressed genes. In stark contrast, the overlap of differentially expressed genes with lamin A/C bound regions was only 20% in wildtype cells similar to the overlap of non-expressed genes, but increased to over 60% in LAP2α knockout cells, including both up-and downregulated genes, which is higher than the overlap with non-expressed genes (Fig. 4A). Thus, lamin A/C may generally spread to gene-rich regions upon LAP2α knockout, but seems to be enriched particularly in regions containing differentially expressed genes. Consistent with these findings, we observed that over 90% of differentially expressed genes overlapped with or were within 10kb distance to the LAP2α knockout–unique lamin A/C peaks - the genomic regions that gained lamin A/C binding in LAP2α knockout versus wildtype cells. This overlap was higher than that with non-expressed genes (Fig. 4B). While the majority of differentially expressed genes in LAP2α knockout versus wildtype cells were located in genomic regions that gained lamin A/C binding in the knockout cells, only less than 30% of these genes overlapped with regions where lamin A/C binding was lost upon LAP2α depletion (Fig. 4B). We concluded that lamin A/C spreads to genomic regions containing most of the deregulated genes in LAP2α knockout versus wildtype cells. To test whether lamin A/C may affect the expression of these genes by binding to the promoter of the genes, we performed heatmap analyses of the lamin A/C ChIP signal around the transcription start sites (TSS). Neither LAP2α nor lamin A/C were enriched at the TSS of differentially expressed genes in both LAP2α knockout and wildtype cells in both the 16 and 20 sonication cycles ChIP samples. In fact, there was even a depletion of these proteins at the TSS compared to surrounding regions, particularly in the 16 cycles ChIP samples (Fig. 4C, Additional file 1: Fig.S4B). In contrast, BRG1 and CHD4 displayed a clear increase in signal on the TSS of both down-and upregulated genes in LAP2α wildtype versus wildtype cells, while their binding to non-expressed genes remained low (Fig. 4C, Additional file 1: Fig.S4B).

**Figure 4.**
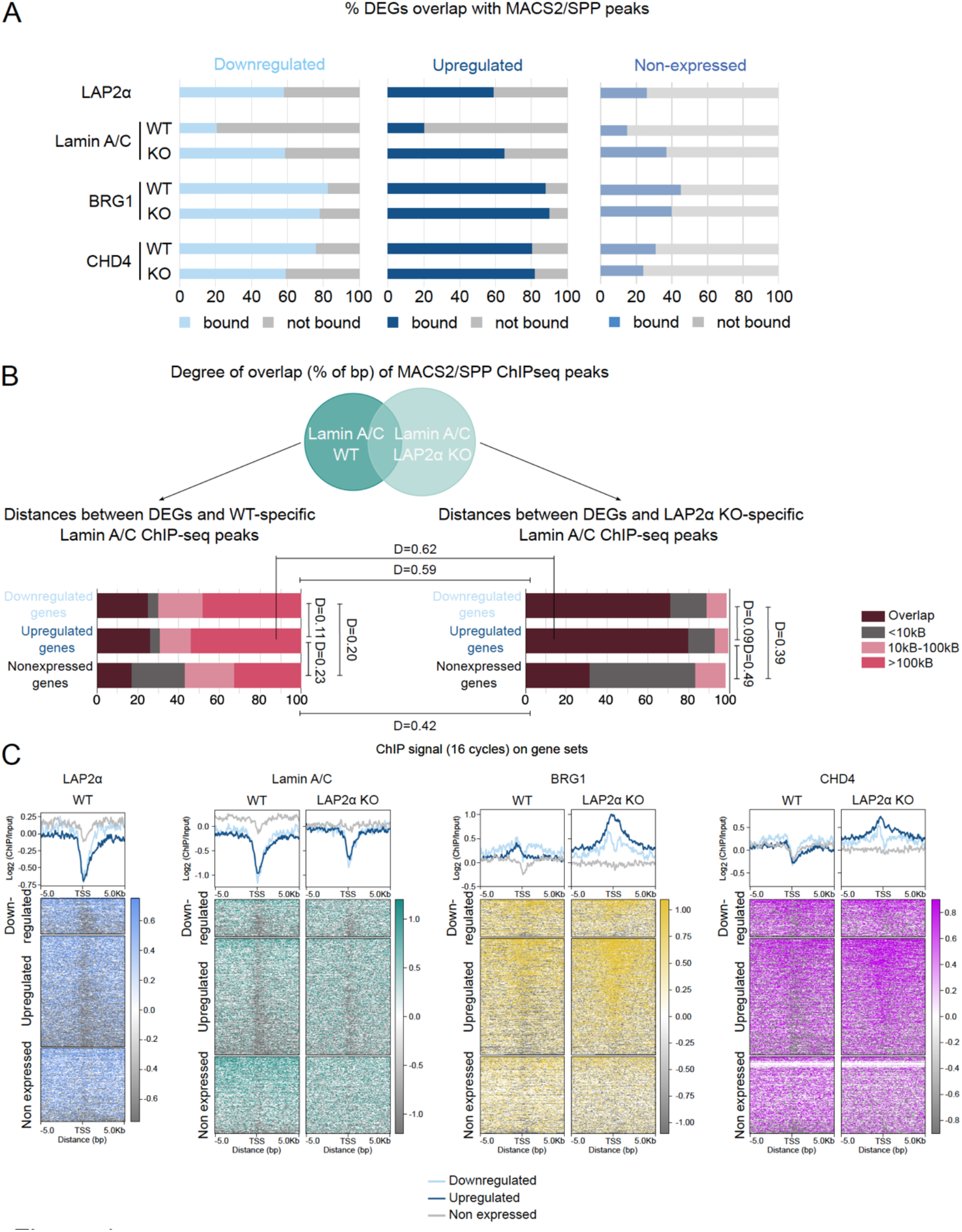
Changes in gene expression correlate with changes in chromatin association of lamin A/C and chromatin remodelers. **(A)** Bar graphs showing the percentages of differentially expressed genes (downregulated – left panel, upregulated – right panel) and 644 non-expressed genes overlapping with combined (16 and 20 sonication cycles) ChIP-seq peaks of LAP2α in wildtype cells and lamin A/C, BRG1 and CHD4 in both LAP2α wildtype and knockout cells. **(B)** Venn diagrams showing the percentage of basepair overlap (% of bp) for combined ChIP-seq peaks of lamin A/C in wildtype and LAP2α knockout cells (upper panel). Bar charts (lower panel) showing genomic distances in base pairs between differentially expressed genes (split into down-and upregulated) and the closest lamin A/C ChIP-seq peaks, subdivided into peaks that are lost upon LAP2α depletion (left panel) and those that are gained (right panel). Distances of non-expressed genes to ChIP-seq peaks are included as a control. Differences between distributions are indicated by the D-value from the KS test, representing the maximum distance between the cumulative distributions of the two sets. All KS tests have an adjusted P-value (P adj) below 0.05. **(C)** Heatmaps showing log2 ratio signals (ChIP over input, 16 sonication cycles) on downregulated (light blue), upregulated (dark blue) and non-expressed (gray) genes +/-5kb from transcription start site (TSS) in LAP2α knockout versus wildtype cells. Log2 ratio signals are presented in the following order from left to right: LAP2α, lamin A/C, BRG1 and CHD4. Graphs above the heatmaps display the mean log2 ratio signals.

In summary, chromatin remodelers BRG1 and CHD4 directly bind to genomic regions with differential accessibility and to differentially expressed genes and their redistribution upon LAP2α depletion correlates directly with changes in both chromatin accessibility and gene expression. The most prominent changes upon LAP2α depletion, however, were observed for lamin A/C, which spreads to open chromatin, containing the majority of deregulated genes and to a lower extent also differentially accessible genomic regions. Interestingly, while lamin A/C spreads to genomic regions around differentially expressed genes, it does not accumulate at the promoters of these genes. Overall, these findings suggest that LAP2α loss causes changes in chromatin accessibility and gene expression, likely mediated in part by chromatin remodelers that directly bind these regions, while the effect of lamin A/C on gene expression and chromatin accessibility seems to be more indirect, probably by affecting chromatin organization genome-wide.

### LAP2α loss causes chromatin reorganization particularly at LAP2α-bound genomic regions

The above results indicate that the loss of LAP2α leads to a gross redistribution of lamin A/C on chromatin, resulting in changes in accessibility and gene expression on a genome-wide level. To confirm these potential changes in chromatin organization through a general and unsupervised analysis, we performed hidden Markov model-based (HMM; [27]) clustering of 5kb genomic bins in LAP2α wildtype and knockout samples separately. The algorithm assigns each bin a, so called hidden state or cluster based on its signals of the obtained genome-wide datasets of the corresponding condition, including RNA-seq, ATAC-seq and ChIP-seq for remodelers and lamin A/C (2 different antibodies 3A6 and E1 were used for lamin A/C ChIP) for 16 and 20 sonication cycles ChIP preparations. To ensure comparability of the assigned clusters in wildtype and knockout samples, we subsequently employed a similarity-based matching algorithm (computed from the respective dataset signals) to arrive at a one-to-one correspondence between wildtype and knockout clusters (see methods for detail). This analysis revealed seven genomic clusters (c1 - c7) with different properties (Additional file 1: Fig. S5A). Following signal enrichment analyses by calculating the log2 fold enrichment of the specific signal in a cluster over the mean genome-wide signal, we grouped the clusters into different functional genomic categories. Clusters c1 to c3 represent active genomic regions based on high enrichment of RNA-seq and ATAC-seq signals, along with an enrichment of remodelers, particularly in clusters c2 and c3 and a depletion of lamin A/C (Additional file 1: Fig. S5A, B). Clusters c4 and c5 exhibit mostly depletion for signals in all datasets, suggesting they may represent highly inactive regions (Additional file 1: Fig. S5A, B). Clusters c6 and c7 represent LAD-type heterochromatic regions because they show low gene expression and chromatin accessibility signal depletion, along with high signal enrichment of A-type lamins (Additional file 1: Fig. S5A, B). The wildtype LAP2α ChIP-seq dataset was not included for clustering to ensure comparability, as LAP2α data was not available for the knockout condition. However, signal enrichment for LAP2α was subsequently calculated within the obtained clusters. LAP2α is slightly enriched in clusters c2 and c5 and prominently enriched in clusters c6 and c7 (Additional file 1: Fig. S5C), which is in line with the known broad distribution of LAP2α including in both active and inactive regions. Although this genome-wide clustering assigned 5kb genomic bins to seven different clusters based on different chromatin properties in both LAP2α wildtype and knockout samples, it did not reveal big changes between knockout versus wildtype samples neither in the assignment of 5kb genomic bins to the 7 clusters (Additional file 1: Fig. S5A, middle panel) nor in the overall signal enrichment values and cluster properties (Additional file 1: Fig. S5B). Similarly, Uniform Manifold Approximation and Projection (UMAP) plots of these genomic clusters did not reveal gross changes between LAP2α wildtype and knockout datasets and no notable changes were detected in cluster similarity to one another (Additional file 1: Fig. S6A). Nevertheless, UMAP plots confirm our observation that lamin A/C is primarily enriched in clusters c6 and c7 while remodelers are enriched in clusters c2 and c3 in both 16 and 20 sonication cycles samples.

As whole-genome clustering analyses revealed only subtle changes, we hypothesized that LAP2α-bound genomic regions in wildtype cells might undergo more dramatic changes upon LAP2α knockout. Therefore, we defined the 5 kb genomic bins with an average LAP2α ChIP log2 signal of 0.6 or greater as LAP2α-bound. These bins covered around 4% of the total genome, which is within the range seen for the genomic coverage of LAP2α, lamin A/C, BRG1 and CHD4 MACS/SPP peaks (see Additional file 2: Table S1). Importantly, these LAP2α-bound 5Kb genomic bins included 84% of downregulated and 53% of upregulated genes upon LAP2α knockout, while only 8% and 3% of chromatin regions with decreased or increased accessibility, respectively (Fig. 5A). This agrees well with the observed overlap of differentially expressed genes and differentially accessible regions with the LAP2α-bound genomic regions identified by peak calling (see Fig. 3 and 4).

**Figure 5.**
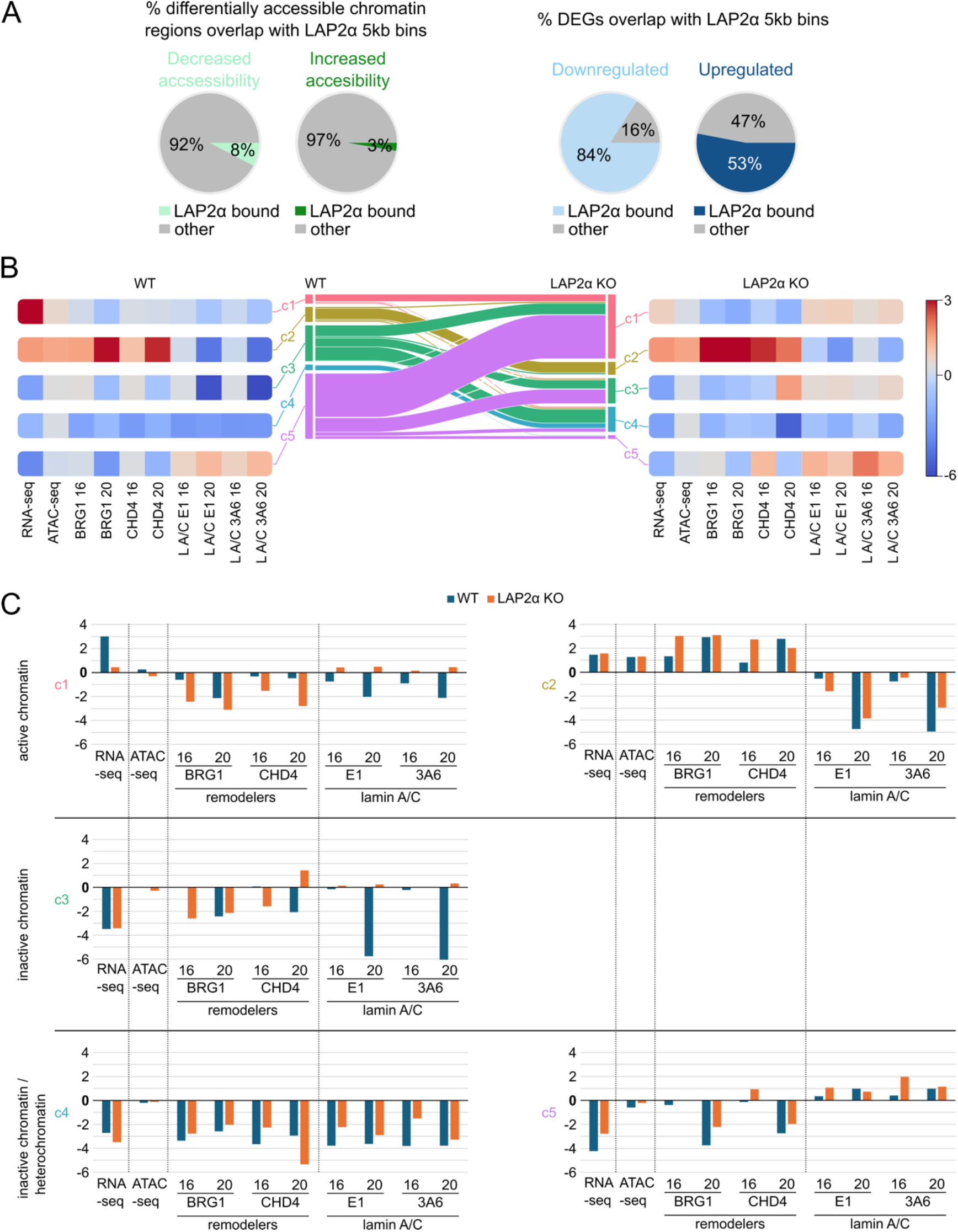
Loss of LAP2α causes chromatin reorganization of LAP2α bound genomic regions. **(A)** Pie charts showing the percentage of differentially accessible regions (left panel) or differentially expressed genes (DEGs; right panel) in LAP2α knockout versus wildtype fibroblasts that overlap with 5 kb genomic bins with a significant LAP2α ChIP-seq signal (minimum log2 fold change of 0.6). **(B)** LAP2α-enriched 5 kb bins were clustered using the Louvain algorithm to compute communities. The middle panel displays a plot showing cluster correspondences between wildtype (WT) and LAP2α knockout (KO) cells. The left panel (WT) and right panel (LAP2α KO) present heatmaps showing signal enrichment for individual parameters, including RNA-seq, ATAC-seq and ChIP-seq (BRG1, CHD4, lamin A/C using E1 and 3A6 antibodies, with 16 and 20 indicating the number of sonication cycles for each sample), relative to the signal in other clusters (log_2_ fold). **(C)** Bar charts displaying signal enrichments from the heatmaps (B) in wildtype and LAP2α knockout cells across all five clusters. The cluster number is indicated on the left side of each chart.

To gain more insight in a possible redistribution of the LAP2α-bound bins defined above, we again employed our clustering and alignment pipeline. To avoid possible problem induced by violation of the assumptions of the HMM-based clustering, we used a graph-based approach to define clusters of LAP2α-bound bins. To this end, we used Louvain community detection on the UMAP graph computed from the signal intensities of all genome-wide datasets in each condition respectively. Subsequent alignment of clusters between wildtype and knockout samples was then performed as before (see Methods for details). This approach identified five clusters which we categorized into active and inactive genomic regions as before (Fig. 5B). In wildtype cells, clusters c1 and c2 show the highest level of gene expression (defined by high RNA-seq signal enrichment) and cluster c2 additionally shows high signal enrichment for chromatin accessibility and chromatin remodelers BRG1 and CHD4 (Fig. 5B, C, WT), indicating that genomic bins in clusters c1 and c2 represent open and active chromatin. Cluster c3 has moderate to no enrichment of these signals, while cluster c4 exhibits the lowest signal enrichments across all parameters (Fig. 5B, C, WT). Cluster c5 shows high enrichment for lamin A/C and low enrichment for all other parameters (Fig. 5B, WT). Thus, clusters c3-c5 likely represent different types of inactive chromatin, with cluster c5 representing mostly heterochromatic LADs. To further characterize these 5 clusters in wildtype cells, we analyzed other properties not used for the clustering algorithm. As expected, clusters c1 and c2 show high overlap with euchromatic ciLADs, while clusters c3 and c4 and particularly c5 show a higher overlap with heterochromatic cLADs (Fig. 6A, WT). Clusters c1, c3 and c4 have similar levels of active (H3K4me3 and H3K9ac) and inactive (H3K27me3 and H3K9me3) histone marks (Fig. 6B, WT), while cluster c2 has higher levels of active histone marks and cluster c5 exhibits higher levels of the inactive H3K9me3 histone mark (Fig. 6B, WT). Furthermore, clusters c1 and c2 have a higher representation of gene-related features, while clusters c3 to c5 are enriched for distal intergenic regions (Fig. 6C, WT).

**Figure 6.**
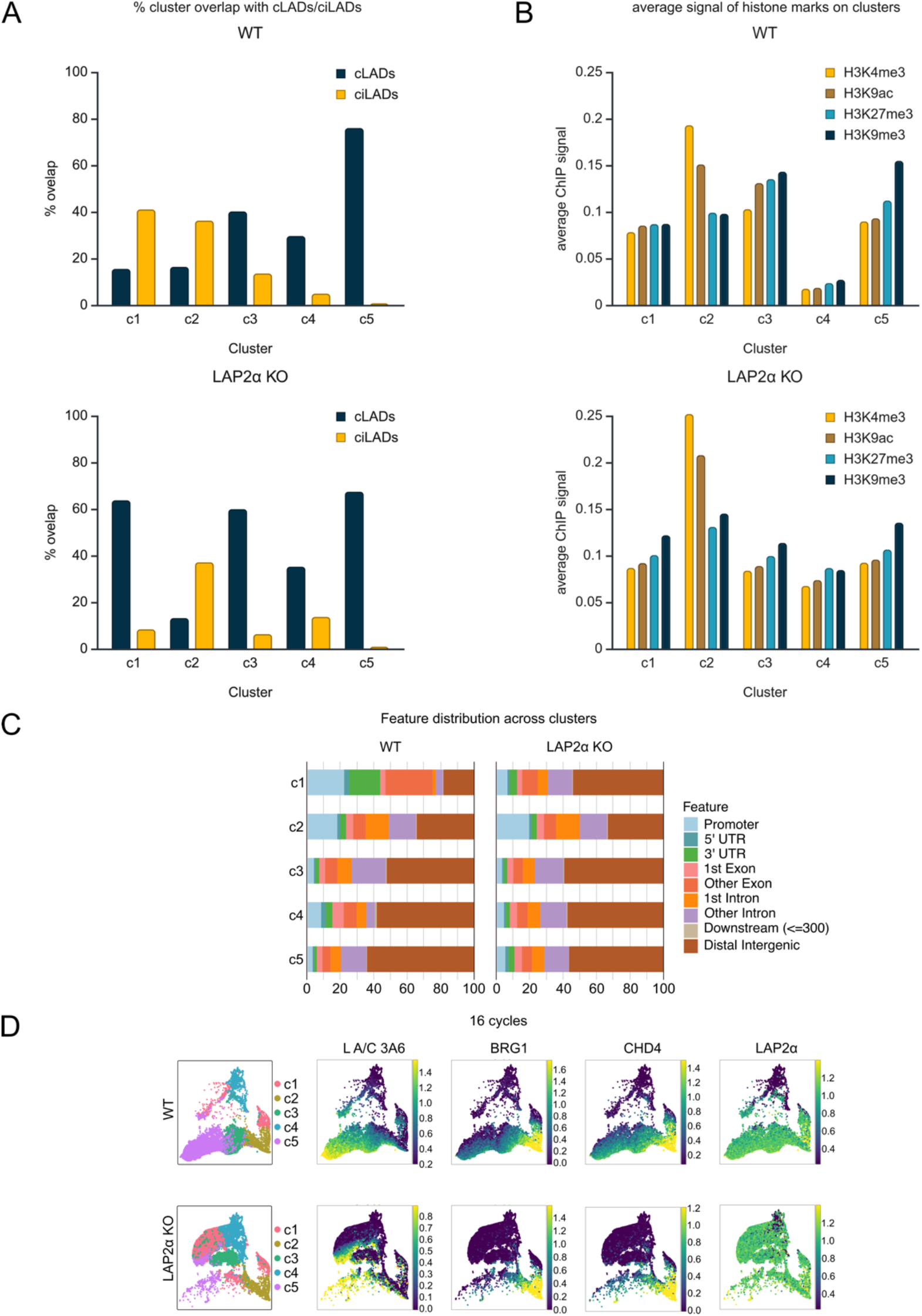
Loss of LAP2α leads to changes in the properties of genomic clusters in LAP2α-bound genomic regions. **(A)** Bar graphs showing the percentages of 5KB bins from each cluster that overlap with constitutive lamina-associated domains (cLADs) and constitutive inter-lamina-associated domains (ciLADs) in wildtype (upper panel) and LAP2α knockout (lower panel) cells. **(B)** Bar graphs showing average ChIP-seq signals of histone marks (H3K4me3, H3K9ac, H3K27me3, H3K9me3) on cluster bins in wildtype (upper panel) and LAP2α knockout (lower panel) cells. **(C)** Bar charts showing the percentages of indicated features overlapping with bins of individual clusters in wildtype (left panel) and LAP2α knockout (right panel) cells. **(D)** LAP2α-enriched bins were clustered by computing communities using the Louvain algorithm on the UMAP graph. The UMAP graph is shown for wildtype (upper panel) and LAP2α knockout (lower panel) cells on the left. On the right, heatmaps plotted on the UMAP graphs display signal intensities for lamin A/C, BRG1, CHD4 and LAP2α ChIP-seq signals (max(ChIP – Input RPM, 0)), starting from left to right.

The differences and similarities of clusters are also visible in the UMAP plots (Fig. 6D, 16 cycles sample, Additional file 1: Fig.S6B, 20 sonication cycles sample). Based on the spatial distribution and proximity of the 5kb bins within clusters in the UMAP plots, one group of genomic bins in cluster c1 seems to resemble bins in cluster 2 (active chromatin), whereas another group of bins within c1 resembles clusters c4 bins (inactive chromatin) in wildtype cells (Fig. 6D). Cluster c3 likely represents a transitional cluster between c2 (active chromatin) and c5 (heterochromatin), being positioned between these clusters on the UMAP (Fig. 6D). As observed in the enrichment plots for the clusters, the UMAPs also show that A-type lamins are primarily localized in cluster c5, while remodelers BRG1 and CHD4 are enriched in cluster c2. In contrast, the plotted signal of LAP2α, analyzed after clustering, is particularly enriched in the active cluster c2, the transitional cluster c3 and the heterochromatic cluster c5 (Fig. 6D, Additional file 1: Fig.S6B), confirming the broad distribution of LAP2α on the genome.

Strikingly, we observed significant rearrangement of the 5 clusters in LAP2α knockout samples (Fig. 5B). Most prominent changes were detected in the LAP2α-bound 5kb genomic bins in wildtype clusters c3 and c5. In particular, 67% of 5kb bins within wildtype cluster c5 are assigned to cluster c1 in LAP2α knockout samples, another group of wildtype cluster c5 bins (22%) to knockout cluster c3 and only a minor group of bins (5%) remained in knockout cluster c5 (Fig. 7A). Similarly, 33% of 5kb bins within wildtype cluster 3 are reassigned to knockout cluster c1, 41% to knockout cluster c4, while the rest remains in knockout cluster c3 (Fig. 7A). This gross reassignment of 5kb bins to different clusters in knockout versus wildtype samples indicates significant changes in the properties of these genomic bins and/or clusters in LAP2α knockout versus wildtype samples. Indeed, the heatmaps showing signal enrichments of lamins and BRG1 and CHD4 in clusters c1 – c5 in LAP2α knockout versus wildtype cells (Fig. 5B), as well as the respective changes in the log_2_ fold signal enrichments (Fig. 5C), reveal that lamin A/C spreads over clusters c1, c3 and c5 in knockout samples, while it is mostly restricted to cluster c5 in the wildtype sample. In contrast, chromatin remodelers BRG1 and to some extent also CHD4 seem to become more enriched in cluster c2 in knockout versus wildtype samples. The spreading of lamin A/C to clusters c1 and c3 and the restriction of BRG1/CHD4 in cluster c2 in the LAP2α knockout cells correlates with reduced gene expression shown by the reduced signal enrichment for RNA, decreased chromatin accessibility and a reduced enrichment of chromatin remodelers particularly in LAP2α knockout cluster c1 (Fig. 5B, C). These changes within the cluster in LAP2α knockout versus wildtype cells are also consistent with an overall increase in cLADs and a decrease in ciLADs in knockout cluster c1 compared to wildtype cluster c1, as well as an increased signal enrichment for the repressive histone mark H3K9me3, an increased overlap with distal intergenic genomic regions and a reduced overlap with gene-related regions (Fig. 6 A-C).

**Figure 7.**
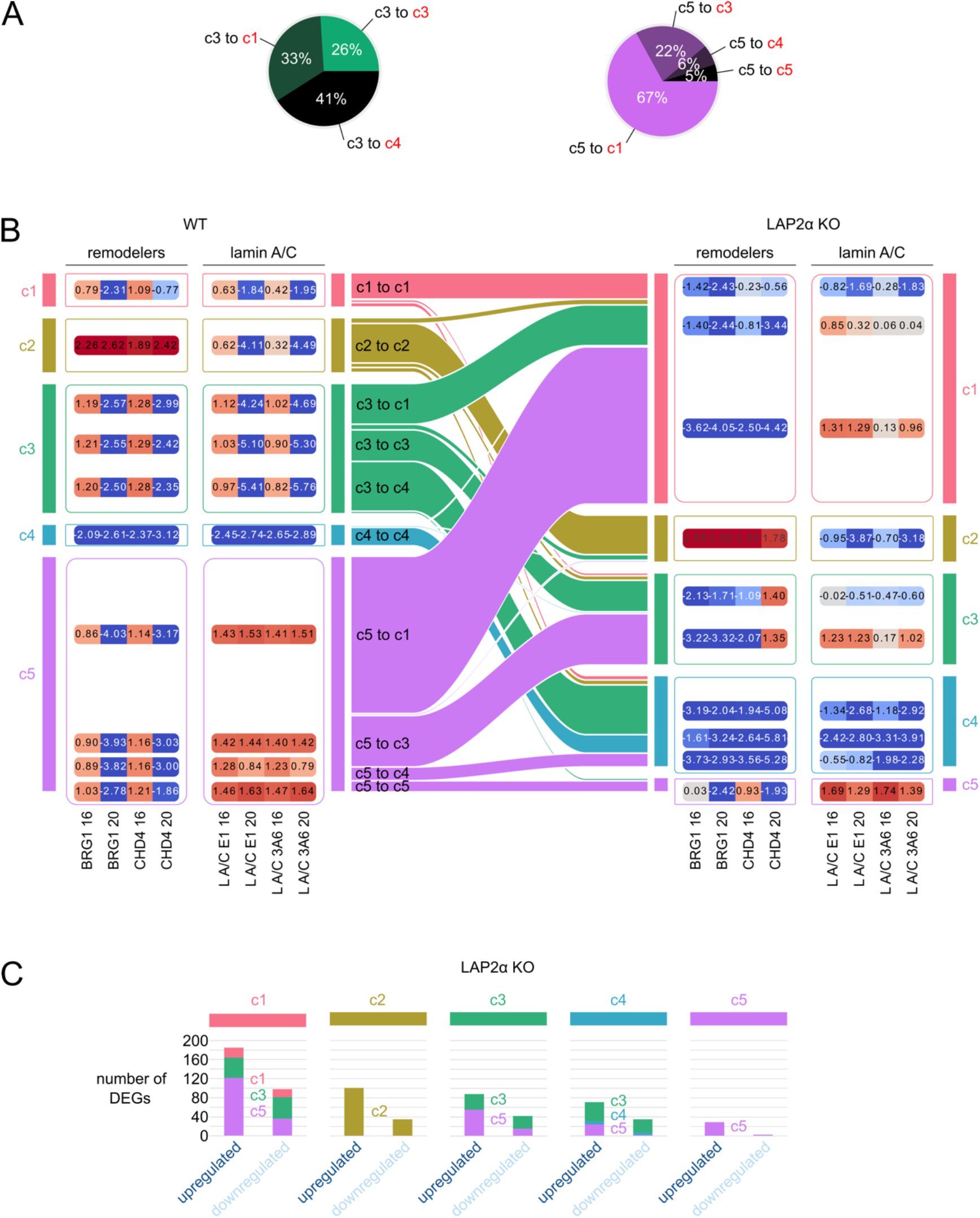
LAP2α depletion causes reorganization of lamin A/C and chromatin remodelers on chromatin, correlating with gene expression changes. Genomic bins forming each cluster were divided into subclusters based on cluster membership in LAP2α wildtype and knockout conditions. **(A)** Pie charts represent the percentage of all bins in wildtype cluster c3 (left) and wildtype cluster c5 (right) that belong to each subclusters switching to other clusters in LAP2α knockout samples (knockout clusters indicated in red). **(B)** Graph showing signal enrichments of chromatin remodelers and lamin A/C across subclusters. The middle panel displays a plot illustrating cluster correspondences between LAP2α wildtype and knockout cells and the subclusters. The left panel (WT) and right panel (LAP2α KO) present heatmaps showing signal enrichment in subclusters for ChIP-seq signals of chromatin remodelers (BRG1 and CHD4) and lamin A/C (immunoprecipitated using E1 and 3A6 antibodies), with 16 and 20 indicating the number of sonication cycles for each sample, relative to the signal in other subclusters (log2 fold). **(C)** Bar graphs showing the numbers of differentially expressed genes (DEGs), divided into upregulated and downregulated genes, overlapping with each cluster in LAP2α knockout (KO) cells. The graphs indicate the numbers of DEGs overlapping with each subcluster that forms the respective clusters in wildtype cells.

The gross reorganization of clusters in LAP2α knockout versus wildtype cells is also evident in UMAP plots. Notably, cluster c1 is massively enlarged and one group of genomic bins in knockout cluster c1 is also positioned between knockout clusters c2 and c5, unlike in the wildtype sample, where cluster c3 is located between clusters c2 and c5. Knockout cluster c3 becomes more similar to clusters c1 and c4 (Fig. 6D). Additionally, the lamin A/C signal, which was mostly found in cluster c5 in wildtype samples, spreads also to clusters c1 and c3 in LAP2α knockout cells (Fig. 5C, 6D and Additional file 1: Fig.S6B). In contrast, BRG1 and CHD4 signals tend to be more efficiently restricted to cluster c2 in knockout versus wildtype cells (Fig. 6D, Additional file 1: Fig.S6B), as already observed in the heatmaps showing signal enrichment for these proteins in the respective clusters (Fig. 5B).

To gain further insights into the redistribution of lamin A/C and remodelers in LAP2α knockout versus wildtype cells, we divided the genomic bins of each cluster into subclusters based on their cluster membership in LAP2α knockout and wildtype samples. The analyses of signal enrichments for these proteins in these defined subclusters revealed that the increased enrichment of lamin A/C in knockout cluster c1 originates mainly from wildtype clusters c3 and c5 (Fig. 7B). Interestingly the genomic bins in these subclusters (c3 to c1 and c5 to c1) show a reduced signal enrichment for remodelers alongside the increased lamin A/C signal enrichment. In contrast, we observed an accumulation of the signals for remodelers exclusively in knockout cluster c2, going hand in hand with a significant reduction of the remodeler signal in all other subclusters in LAP2α knockout versus wildtype samples (Fig. 7B). The observed increase of lamin A/C signal enrichment in knockout cluster c3 (Fig. 5B) originates mostly from genomic bins within wildtype cluster c5 moving to c3 in LAP2α knockout sample (Fig. 7B). In contrast, knockout cluster c4, despite gaining genomic bins mostly from wildtype clusters c3 and c5, retains its depletion of both lamins and remodelers (Fig. 7B). Knockout cluster c5 becomes much smaller in LAP2α knockout versus wildtype cells but remains high in lamin A/C enrichment (Fig. 7B).

Interestingly the observed changes in cluster assignment of genomic bins in LAP2α knockout versus wildtype samples correlate well with alterations in gene expression. The highest number of differentially expressed genes (DEGs) are found in the most significantly rearranged cluster in the knockout sample (c1), the majority of which originates from wildtype cluster 5 (Fig. 7C). Similarly, knockout cluster c2, which shows a significant enrichment for remodelers, contains a large number of mostly upregulated genes (Fig. 7C).

Overall, unsupervised clustering of LAP2α-bound 5kb genomic bins revealed a gross rearrangement of these genomic regions in LAP2α knockout versus wildtype cells leading to a spreading of lamins to active chromatin regions, accompanied by a restriction of nucleosome remodelers to a subset of active genomic regions and a reduction in gene expression and chromatin accessibility in lamin A/C-bound regions.

## Discussion

This study examines the genome-wide changes in chromatin organization in mouse dermal fibroblasts following LAP2α depletion. We demonstrate that loss of LAP2α leads to alterations in chromatin accessibility and gene expression, correlating with A-type lamin spreading to active genomic regions. Lamin spreading is also accompanied by a redistribution of nucleosome remodelers on chromatin and their enrichment on highly active genomic regions, directly correlating with changes in chromatin accessibility and gene expression genome-wide.

LAP2α is a well-known regulator of the nucleoplasmic pool of A-type lamins. Our previous studies have revealed some mechanistic insight into the effect of LAP2α on lamin A/C chromatin association, showing that loss of LAP2α leads to a stronger and/or more stable binding of lamin A/C to chromatin [12, 13]. Our working model proposes that in wildtype cells, LAP2α may restrict lamin A/C chromatin association by competing for chromatin binding sites and/or by forming non-chromatin bound nucleoplasmic complexes with lamin A/C and BAF [12]. The regulation of lamin A/C chromatin association by LAP2α may play critical roles in tissue homeostasis by affecting the transition of tissue progenitor cells from a quiescent to a proliferative or differentiating state [16, 18, 19]. For example, LAP2α loss in mice delays myogenic differentiation, likely through a deregulation of lamin A/C chromatin association allowing lamin A/C binding to active genomic regions and impairing myogenic gene regulation[16, 18].

Although our previous findings supported a role of LAP2α in chromatin organization through the regulation of lamin A/C, it remained unclear whether LAP2α depletion may also affect other major chromatin regulating proteins thereby contributing to the observed phenotypes in LAP2α knockout cells. To address this important open question and to obtain a broader perspective on genome-wide changes following LAP2α loss, we included in our study here the ATPase subunits of the nucleosomal remodelers SWI/SNF and NURD complexes, BRG1 and CHD4, respectively. Chromatin remodelers are ATP-dependent enzymes that regulate nucleosome positioning and composition, controlling DNA accessibility for transcription, replication and repair [22]. Their activity is tightly regulated through histone modifications, DNA sequences and interactions with transcription factors, ensuring precise chromatin organization [28]. Remodelers localize to key genomic regions such as promoters, enhancers and replication origins, where they establish phased nucleosome arrays downstream of transcription start sites, facilitating transcription elongation while ensuring stable chromatin organization in gene bodies [29, 30]. Additionally, certain remodelers like SWI/SNF are involved in histone variant exchange, which plays a role in transcription start site selection and chromatin stability [31, 32]. Their dynamic “hit-and-run” mechanism of action allows transient chromatin interactions and rapid adaptability [33, 34]. Importantly, components of these complexes have been found in previous studies to associate with lamin complexes in various contexts [20, 35] and our previous study on the analyses of the LAP2α-interactome in a proximity-based assay indicated that LAP2α may also interact with BRG1 and CHD4 containing complexes [23]. We confirmed here by co-immunoprecipitation analyses that BRG1 and CHD4 may indeed form complexes with LAP2α and lamin A/C and that binding of BRG1 and CHD4 to lamins occurs independently of LAP2α and vice versa.

Interestingly, we found that LAP2α knockout in mouse fibroblasts not only leads to altered chromatin binding of lamin A/C, but also to a redistribution of BRG1 and CHD4 on chromatin to some extent, which correlated with changes in chromatin accessibility and gene expression. Our results revealed that in wildtype conditions, genomic regions enriched for LAP2α and for chromatin remodelers are often in close proximity or overlap with each other, whereas lamin A/C-bound regions are spatially separated from BRG1-and CHD4-enriched sites on chromatin. Upon LAP2α depletion, lamin A/C relocates towards open genomic regions closer to sites enriched for BRG1 and CHD4 and around differentially expressed genes, correlating with changes in chromatin accessibility. As BRG1 and CHD4, but not lamin A/C, bind to genomic regions showing differential accessibility and to deregulated genes in LAP2α knockout versus wildtype cells, we propose that the redistribution of chromatin remodelers, rather than the changes of lamin A/C, are the direct mediators of changes in chromatin accessibility and gene expression. Unsupervised clustering of LAP2α-enriched 5 kb genomic bins based on signal enrichments across all datasets, including RNA-seq, ATAC-seq and lamin A/C, BRG1 and CHD4 chromatin immunoprecipitation data revealed that LAP2α depletion causes genome-wide reorganization of chromatin. Additionally, this clustering approach confirmed spreading of lamin A/C to active genomic regions upon LAP2α loss correlating with decreased gene expression. Interestingly, lamin A/C spreading was linked to enrichment of chromatin remodelers in a subset of highly active chromatin clusters, while they were reduced to some extent in other active genomic regions. Notably, the majority of differentially expressed genes in knockout cells (84% of downregulated and 53% of upregulated genes) were located in LAP2α-bound regions in wildtype cells. A large proportion of these genes either overlapped with regions that gained lamin A/C binding in LAP2α knockout cells or were enriched in the clusters of genomic bins showing enrichment for chromatin remodelers following LAP2α loss.

While this study offers novel insights into the previously observed effects of LAP2α depletion on chromatin organization, it also raises several new questions. It is still unclear, whether the redistributions of lamin A/C and chromatin remodelers on chromatin upon LAP2α depletion are independent or directly linked events and what are primary and direct effects of the LAP2α knockout. The fact that both LAP2α and lamins A/C can be found in nuclear complexes of BRG1 and CHD4 individually or together may argue for some coordination of these redistributions. However, the molecular mechanisms how lamins can affect BRG1 and CHD4 or vice versa remain elusive. This topic is also linked to another major open question on how lamin spreading to open chromatin in LAP2α knockout cells affects gene expression and chromatin accessibility, both of which are directly correlated with changes in nucleosomal remodeler binding. As lamin A/C does not enrich directly on genes or on genomic regions showing altered accessibility in LAP2α knockout versus wildtype conditions, one can exclude direct competition of binding between lamin A/C and chromatin remodelers. One possibility is that lamin A/C may indirectly affect these processes by forming large compartments on open, gene-rich genomic regions, which may recruit or exclude specific transcription and chromatin regulators, including remodelers. Such a scenario is supported by our previous findings that in the absence of LAP2α, lamin A/C forms large immobile lamin A complexes that seem to interact more strongly and stably with chromatin [8, 12]. This hypothesis directly leads to the question as to how LAP2α can prevent lamin A/C from forming these stable complexes on chromatin. A recent study suggests that LAP2α can affect chromatin binding of lamin A/C by at least two non-mutually exclusive mechanisms, by competing with lamin A/C for chromatin binding and by recruiting lamin A/C to soluble, non-chromatin-bound complexes in the nucleoplasm [12].

Overall, based on the results shown in this study together with previous observations, we propose that LAP2α restricts A-type lamin binding to active chromatin regions under wildtype conditions, allowing chromatin remodelers to modulate chromatin accessibility, thereby contributing to gene regulation. In the absence of LAP2α, lamin A/C spreads to active genomic regions, where it can displace or restrict access of chromatin remodelers and possibly other proteins, which in turn leads to chromatin reorganization, altered accessibility and abnormal gene expression.

## Conclusions

Our study highlights the critical role of LAP2α in chromatin organization and regulation. LAP2α associates with chromatin and with a dynamic pool of lamin A/C in the nuclear interior in a complex, inter-dependent manner [12] and thereby may prevent lamin A/C from forming stable complexes on open gene-rich genomic regions, enabling chromatin remodelers and other chromatin regulators to access these genomic regions to regulate accessibility and gene expression. We hypothesize that LAP2α depletion leads to lamin A/C spreading into active chromatin regions, which in turn causes global changes in chromatin organization. The lamin A/C mediated chromatin reorganization is correlated with redistribution of chromatin remodelers on chromatin, directly affecting chromatin accessibility and gene expression. These findings highlight the general importance of higher order chromatin organization in gene expression besides the more specific regulation of gene transcription. Our study also provides potential mechanisms how the complex interplay between lamins and LAP2α and their chromatin association can indirectly affect chromatin accessibility and gene expression, although at this point we cannot distinguish primary and secondary effects caused by LAP2α knockout.

## Methods

### Cell lines

Mouse dermal fibroblasts, including LAP2α wildtype (WT), LAP2α knockout (KO) and LAP2α/lamin A/C double knockout (DKO) variants, were generated using the CRISPR/Cas9 system as previously described [8]. These cells were cultured in Dulbecco’s modified Eagle’s medium (DMEM) supplemented with 10% fetal calf serum (FCS), 2 mM glutamine, 100 U/ml penicillin, 100 μg/ml streptomycin (P/S) (all from Sigma-Aldrich) and non-essential amino acids (from PAN-Biotech), under humidified conditions at 37°C with 5% CO2.

### Immunoprecipitation and immunoblotting

Co-immunoprecipitation (co-IP) and immunoblotting were performed as described previously [12], with some modifications. In brief, cells were scraped into an IP buffer, lysed on ice and then centrifuged to remove insoluble material. The cleared supernatant was pre-cleared with protein A/G beads, followed by overnight incubation with antibodies while rotating. Protein–antibody complexes were then pulled down, washed and eluted in SDS–PAGE sample buffer for subsequent immunoblot analysis. For immunoblotting, membranes were incubated with the following primary antibodies: anti-BRG1 (1:10,000; abcam: ab110641), anti-CHD4 (1:10,000; abcam: ab70469), anti-lamin A/C E1 (1:1,000; Santa Cruz Biotechnology: sc-376248), anti-FLAG (1:1,000; Sigma Aldrich: F3165), anti-LAP2α 1H11 (1:100; Max Perutz Labs Monoclonal Antibody facility, see also [13]) and anti-γ-tubulin (1:5,000; Sigma Aldrich: T6557). Uncropped images of Western blots are shown in the Additional file 3.

### Chromatin immunoprecipitation sequencing (ChIP-seq)

Chromatin immunoprecipitation was performed as described previously [12], with following modifications: before DNA-protein fixation, a protein-protein fixation step was introduced. Cells were pelleted and resuspended in 2 mM DSG (disuccinimidyl glutarate) in PBS (0.5 ml per 1 million cells) and incubated for 20 minutes at room temperature with gentle rotation. Formaldehyde fixation was then performed as described previously and all subsequent steps were carried out according to the original protocol. In brief, cells were harvested, crosslinked first with DSG and then formaldehyde, followed by quenching with glycine. After washing, cells were lysed and chromatin was fragmented by sonication (16 cycles of 30 seconds ON/30 seconds OFF) using a Bioruptor Pico sonication device (Diagenode). Fragmented chromatin was incubated overnight with the following antibodies for immunoprecipitation: anti-BRG1 (30 μl/ChIP; abcam: ab110641), anti-CHD4 (30 μl/ChIP; abcam: ab70469), anti-lamin A/C E1 (60 μl/ChIP; Santa Cruz Biotechnology: sc-376248), anti-lamin A/C 3A6 (300 μl/ChIP; Max Perutz Labs Monoclonal Antibody facility, see also [13]) and anti-LAP2α 1H11 (300 μl/ChIP; Max Perutz Labs Monoclonal Antibody facility, see also [13]). Immunoprecipitated chromatin was captured using Protein A/G beads. After stringent washing, chromatin was eluted, crosslinks reversed overnight and DNA purified for further analysis. After the standard protocol was completed, half of the input and immunoprecipitated chromatin was further sonicated for an additional 4 cycles of 30 seconds ON/30 seconds OFF. Both chromatin preparations (16-cycle and 20-cycle sonications) were submitted for sequencing.

The DNA for ChIP-sequencing was processed at the Next Generation Sequencing facility of the Vienna Biocenter Core Facilities (VBCF; https://www.viennabiocenter.org/vbcf/next-generation-sequencing/). The library was prepared using the NEBNext® Ultra™ II DNA Library Prep Kit for Illumina and the samples were sequenced on an Illumina NovaSeq SP platform in SR100 mode (single-end reads; 100 bp length).

Raw reads were mapped to the mouse genome assembly *Mus musculus* GRCm38 using NextGenMap [36] v0.5.5 using default settings. The mapped reads were filtered for a minimum mapping quality of 10 using samtools view-q 10 and indexed using samtools index. The log2ratio files between the ChIP and Input samples were generated using bamCompare from deepTools with the settings --operation log2 --binSize 10. Peaks were called using two different programs, with MACS2 [37] v2.1.4 using settings callpeak --nomodel-g mm-p 1e-1 --extsize 200 and with SPP [38] v1.14 with default settings. Peak regions that are common in both MACS2 and SPP peaks were extracted for each sample using bedtools intersect. To optimize these peak intersections for the regions with maximum CHIPseq signal, HOMER [39] v4.11.1 was used, by first running makeTagDirectory on all mapped read files of all samples, then by running getPeakTags-center on the peak files. Resulting peak sets were used to be combined for 16 and 20 sonication cycles of the same samples using bedtools merge.

For lamin A and LAP2α samples, peaks were also called using Enriched Domain Detector (EDD) [24] v1.1.19 with default settings, which were used for data interpretation, but not for downstream analyses.

Reproducibility of the ChIP-seq samples was evaluated by calculating the Pearson correlation coefficients. The mapping coverages of the short sequencing reads of each sample were calculated in 2500 base pair bins across the genome using deepTools multiBamSummary and the correlation between the samples was analyzed using deepTools plotCorrelation with the settings --corMethod pearson -- removeOutliers.

### RNA-sequencing (RNA-seq)

Cells were plated on 10 cm culture dishes 48 hours prior to harvesting. Total RNA was isolated using the RNeasy® Mini Kit (Qiagen) following the manufacturer’s protocol. The extracted RNA was then sent to the Next Generation Sequencing facility at the Vienna Biocenter Core Facilities (VBCF; https://www.viennabiocenter.org/vbcf/next-generation-sequencing/) in Vienna, Austria, for library preparation. This included poly(A) mRNA enrichment using the NEBNext® Poly(A) mRNA Magnetic Isolation Module and subsequent library preparation with the NEBNext® Ultra™ II Directional RNA Library Prep Kit for Illumina (both from New England Biolabs). Sequencing was performed on the Illumina NextSeq 550 platform in PE75 mode (paired-end reads; 75 bp length).

Raw reads were mapped to the reference mouse genome assembly *Mus musculus* GRCm38 using STAR [40] v2.7.10b with default settings using the mouse genome annotation GRCm38.102 from ENSEMBL [41]. Differential expression between LAP2α KO and WT samples was analyzed using Rsubread [42] v2.8.2 and DESeq2 [43] v1.34.0 and the figure representing the expression of the genes was plotted with ggplot2 (https://ggplot2.tidyverse.org) v3.3.5. The R package qvalue (http://github.com/jdstorey/qvalue) was used to calculate the q-values of the p-values at an FDR level of 0.05. Genes with a mean minimum of 5 mapped reads across all replicates, minimum log2 fold change of 2 and a maximum local FDR value of 0.05 were selected as significantly differentially expressed genes. A mean number of genes of those that are up and down regulated were randomly selected among the genes that showed very little to no expression (average of total mapped reads across 6 replicates for each gene = 0.35) and were used as a control in the analyses as the random non-expressed gene set. Mapping files of 3 replicates of each WT and LAP2α KO samples were merged into single files using samtools merge and representative signal track files were generated using bamCoverage of deepTools with RPKM normalization method.

Differentially expressed genes are listed in the Additional file 4: Table Differential Expression.

### ATACseq

Cells were plated on 15 cm culture dishes 48 hours before harvesting and sent to the Next Generation Sequencing facility at the Vienna Biocenter Core Facilities (VBCF; https://www.viennabiocenter.org/vbcf/next-generation-sequencing/) in Vienna, Austria, for protocol execution and sequencing. In brief, cells were harvested and washed prior to nuclei isolation. Nuclei were prepared by gentle lysis, followed immediately by chromatin transposition using Tn5 transposase. After the transposition step, DNA was purified and amplified via PCR. The resulting libraries were purified and prepared for sequencing. Sequencing was carried out on the Illumina NextSeq 550 platform in SR150 mode (single-end reads; 150 bp length). Raw reads were trimmed using Trimmomatic [44] v0.39 with these settings: NexteraPE-PE.fa:2:30:10 MINLEN:36. ATACseq reads were mapped to the reference mouse genome assembly *Mus musculus* GRCm38 using bowtie2 [45] v2.5.1 with the setting --very-sensitive and then were sorted by read name using samtools [46]. ATACseq peaks were called using Genrich (https://github.com/jsh58/Genrich) using 3 replicates of each WT and KO with the settings-j-y-r-e MT-q 0.01.

Differential accessibility between LAP2α KO and WT ATACseq data on the ATACseq peak regions was analyzed using DiffBind [47] v3.4.9 and the genomic regions showing a minimum difference of 1 in log2-transformed values and FDR <0.01 of accessibility signal were defined as differentially accessible regions. A mean amount of these regions with increased or decreased accesibility was randomly selected across the genome using bedtools shuffle in the same lengths and on the same chromosomes as the original peaks were, to be used as the randomized regions as a control for ATACseq analyses.

To be used for single representation of ATACseq data on the figures, the mapped reads of the three replicates of each WT and KO samples were merged using samtools merge and then converted into signal track files using bamCoverage from deepTools using RPKM normalization.

The differentially accessible regions are shown in the Additional file 4: Table Differential Accessibility.

### Additional computational methods

The raw sequencing read qualities were evaluated using FastQC (http://www.bioinformatics.babraham.ac.uk/projects/fastqc/).

The closest distances between genomic regions of interest were calculated using closest-features --dist from BEDOPS [48] v2.4.39 and were illustrated by summarizing these distances in bins of lengths. The randomized peaks in the pairwise closest distance plots (Fig. 2B and S3A) were generated for 100 times using bedtools shuffle for the peak set B that peak set A is being compared with, by randomly selecting regions in the genome in the same lengths and amount as the original peaks maintaining the same distribution across chromosomes and the averages of the closest distances of these 100 randomized peak sets versus the peak set A were illustrated. The significance of the distributions of the closest distances between genomic regions of interest were tested using the Kolmogorov-Smirnov test (KS test) in R Statistical Software (v4.0.3; R Core Team 2020) using dgof package. p.adjust function in R with the Hochberg method was used to correct the p-values of the relevant groups of KS tests.

Signal track files of the mapped reads were generated using bamCoverage from deepTools [49] v3.5.1 using RPKM normalization method. The log2ratio and RPKM coverage files were illustrated as heatmaps with average signals on regions of interest using computeMatrix and plotHeatmap functions of deepTools and using IGV [50]. When required, file formats were converted using bigWigToBedGraph and bedGraphToBigWig [51].

Distributions of differentially accessible regions and differentially expressed genes per chromosome were calculated by normalizing the percentages of distribution per chromosome to the chromosome lengths.

Overlapping peak and gene coordinates between samples of interest were found using bedtools [52] v2.29.2 intersect command.

cLAD and ciLAD regions were downloaded from GEO with the ID GSE17051 for mm9 mouse genome assembly and their genomic coordinates were converted to mm10 assembly coordinates using hgLiftOver from UCSC [53].

The feature distribution figure (Fig. 6C) was generated using ChIPseeker [54] v1.26.2. The signal summary plots in Fig. 1.F were generated using R. bigWigAverageOverBed from KentUtils [55] was used to generate average log2 or RPKM signals on regions of interest.

### Clustering

#### Whole genome chromHMM segmentation

Whole genome and all annotated genes were subdivided into groups of similar signal signatures based on read counts of all available datasets in this study (BRG1-, CHD4-and LaminA/C (3A6)-and-lamin A/C (E1) ChIP-seq, 16 and 20 sonication cycles each, ATAC-seq and RNA-seq). In brief, we segmented the genome using chromHMM v1.24[27] by first binarizing the alignment data into 5kb bins with BinarizeBam (-mixed) and subsequently fitting a 10-state Hidden Markov Model (HMM) with learnModel (-p 8-r 1000-b 100000-printstatebyline).

#### Per bin read quantification

Sequencing data was counted per 5kb bin using deeptools bamCoverage v3.4.3[49]. In brief, we quantified the data with bamCoverage (--ignoreDuplicates-p 8-of bigwig-bs 5000). We then merged the resulting counts into a single table and subsequently normalized it by either computing counts per million reads (CPM) for signal and input data and then subtracting the mean input CPM from the respective signal CPM value per bin using min (difference, 0) to ensure only positive values (ChIP-seq) or simply computing CPM (ATAC-seq, RNA-seq).

#### Louvain clustering of genomic bins

Genomic bins were clustered by computing communities using the Louvain algorithm on the UMAP graph. In brief, we first assigned each bin its corresponding signal vector (quantified as described above) and used this as input for the UMAP algorithm implemented in umap-learn v0.5.3 [56]. Subsequently, we used Louvain community detection with a resolution of 0.15 (implemented in networkx v3.1 [57]) to partition the resulting UMAP graph into distinct clusters. This procedure was used to cluster either the whole genome or LAP2α-enriched bins only. The latter were computed by calculating the average log2 ratio of LAP2α and its input (1H11 WT 16 sonication cycles samples) for 5kb bins of the whole genome using bigWigAverageOverBed from KentUtils and subsequently thresholding at a value of 0.6 where all bins above this value were said to be LAP2α-enriched.

#### Computing cluster correspondences between WT and KO

Cluster correspondences between LAP2α wildtype and knockout data were computed using a PCA-based cluster matching algorithm. In brief, we first computed enrichment vectors for each cluster by dividing the mean log-normalized signal over bins within a cluster by the mean log-normalized signal over all bins:

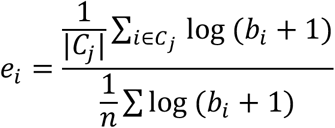

where *C_j_* denotes a cluster *j* of a given condition, *b_i_* denotes the normalized signal of a given sequencing data set in genomic bin *i* and *n* is simply the number of bins involved in this analysis. To counteract the course of dimensionality, we then subjected the computed enrichment vectors to a principal component analysis (PCA, implemented in scikit-learn v1.3.0; https://www.jmlr.org/papers/v12/pedregosa11a.html) and used the first two principal components to compute pairwise L1-distances (implemented in scipy v1.11.2;[58]) between the clusters of wildtype and knockout datasets. We then used the resulting data compute bipartite nearest neighbour graph between wildtype and knockout clusters and subsequently found groups of corresponding clusters by computing the connected components of the resulting graph.

The list of genomic bins and cluster memberships of whole genome clustering and clustering done on LAP2α bound genomic bins are shown in the Additional file 5.

## Supplemental Information

Additional file 1: Supplemental Figures: Fig. S1-S6.

Additional file 2: Supplemental Table S1. Summary of ChIP-seq peaks parameters obtained for different samples and using different algorithms.

Additional file 3: Images of full western blots.

Additional file 4: Tables of differentially expressed genes and differentially accessible regions.

Additional file 5: Tables of cluster memberships of genomic bins in whole genome clustering and LAP2α-bound bin clustering.

## Declarations

### Ethics approval and consent to participate

Not applicable

### Consent for publication

Not applicable

### Availability of data and materials

The RNAseq, ChIPseq, and ATACseq datasets generated during the current study are available in the GEO repository, under these links: (https://www.ncbi.nlm.nih.gov/geo/query/acc.cgi?acc=GSE292284), (https://www.ncbi.nlm.nih.gov/geo/query/acc.cgi?acc=GSE292285), (https://www.ncbi.nlm.nih.gov/geo/query/acc.cgi?acc=GSE292286).

The histone mark ChIPseq data analysed during the current study are available in the GEO repository, under these links:

H3K9ME3.KO (https://www.ncbi.nlm.nih.gov/geo/query/acc.cgi?acc=GSM1717512) H3K9AC.KO (https://www.ncbi.nlm.nih.gov/geo/query/acc.cgi?acc=GSM1717506) H3K4ME3.KO (https://www.ncbi.nlm.nih.gov/geo/query/acc.cgi?acc=GSM1717508) H3K27ME3.KO

(https://www.ncbi.nlm.nih.gov/geo/query/acc.cgi?acc=GSM1717510) H3K27ME3.WT

(https://www.ncbi.nlm.nih.gov/geo/query/acc.cgi?acc=GSM1717509)

H3K4ME3.WT (https://www.ncbi.nlm.nih.gov/geo/query/acc.cgi?acc=GSM1717507)

H3K9AC.WT (https://www.ncbi.nlm.nih.gov/geo/query/acc.cgi?acc=GSM1717505) H3K9ME3.WT (https://www.ncbi.nlm.nih.gov/geo/query/acc.cgi?acc=GSM1717511) The cLAD and ciLAD regions used in this study are available on GEO with accession code GSE17051 (https://www.ncbi.nlm.nih.gov/geo/query/acc.cgi?acc=GSE17051). Source code for in-house scripts that are used for analyses are available on: https://github.com/FatihSarigol/Filipczak_et_al_2025.

### Authors’ contributions

Conceptualization, D.F., R.F. and N.N.; Performance of the experiments, D.F; Bioinformatic analysis, F.S., D.M.; Methodology, D.F., F.S, D.M; Data interpretation and writing the manuscript, D.F., F.S., R.F. and N.N.; Funding Acquisition, R.F and N.N.; Supervision, R.F. and N.N. All authors read and approved the final manuscript.

## Supporting information

Additional file 3_uncropped images of Western blots

Additional file 4_Differential_Accessibility_and_Expression

Additional file 5_Clustering

## Acknowledgements

The NGS sequencing was performed by the Next Generation Sequencing Facility at Vienna BioCenter Core Facilities (VBCF), member of the Vienna BioCenter (VBC), Austria (https://www.viennabiocenter.org/vbcf/next-generation-sequencing/).

---

**Additional file 1**

**Supplemental Figures: Fig. S1-S6**

**Figure S1.**
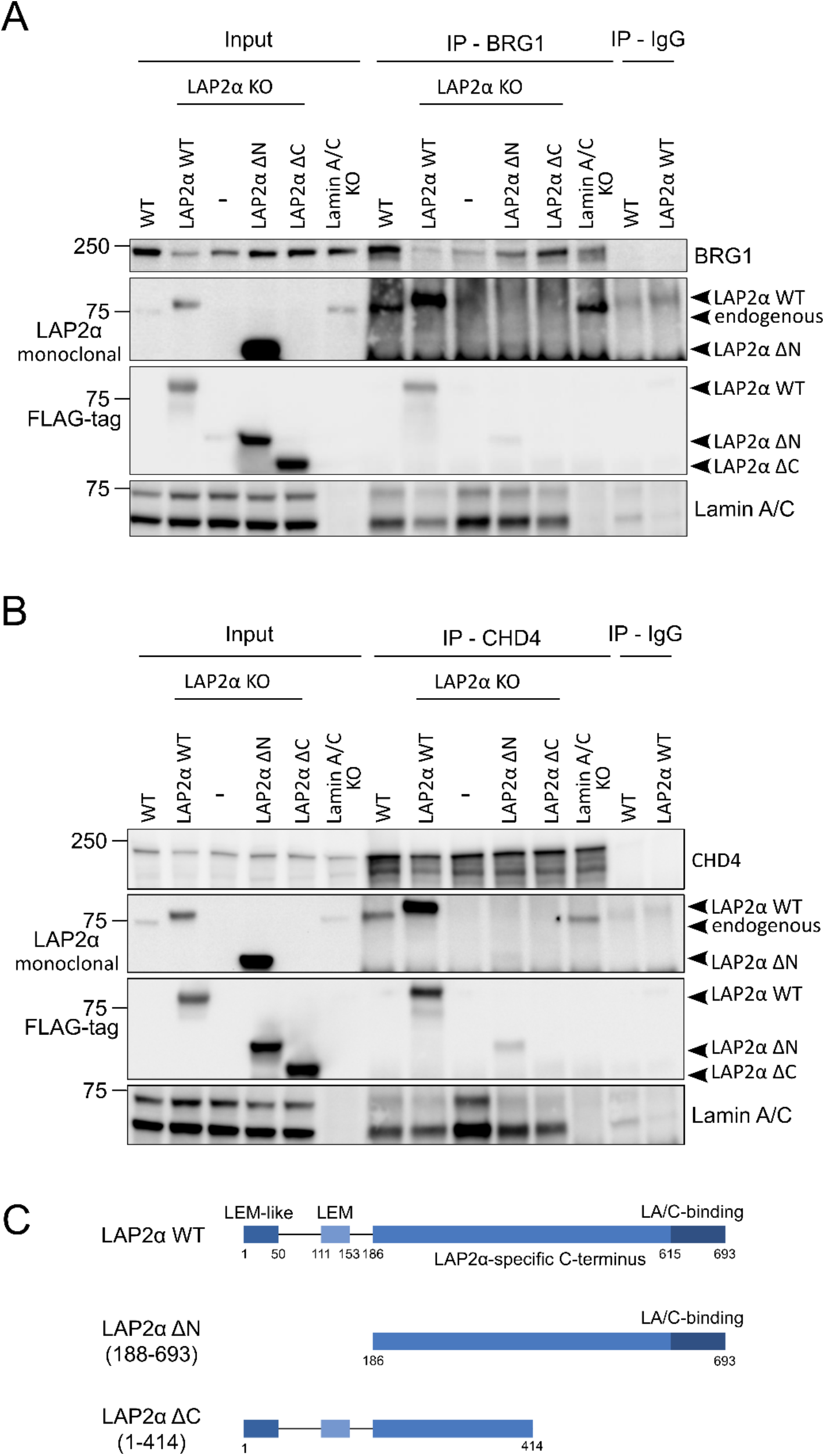
Chromatin remodelers BRG1 and CHD4 coprecipitates with lamin A/C and LAP2α. (A, **B)** BRG1 (A) and CHD4 (B) were immunoprecipitated from wildtype, LAP2α KO, Lamin A/C KO and LAP2α KO cell lines ectopically expressing LAP2α WT, LAP2α ΔN or LAP2α ΔC. The immunoprecipitated samples were subsequently subjected to Western blot analysis using the indicated antibodies. **(C)** Graphical representation of LAP2α and its truncation variants that were expressed in LAP2α knockout (KO) cell lines and used in co-immunoprecipitations assays.

**Figure S2.**
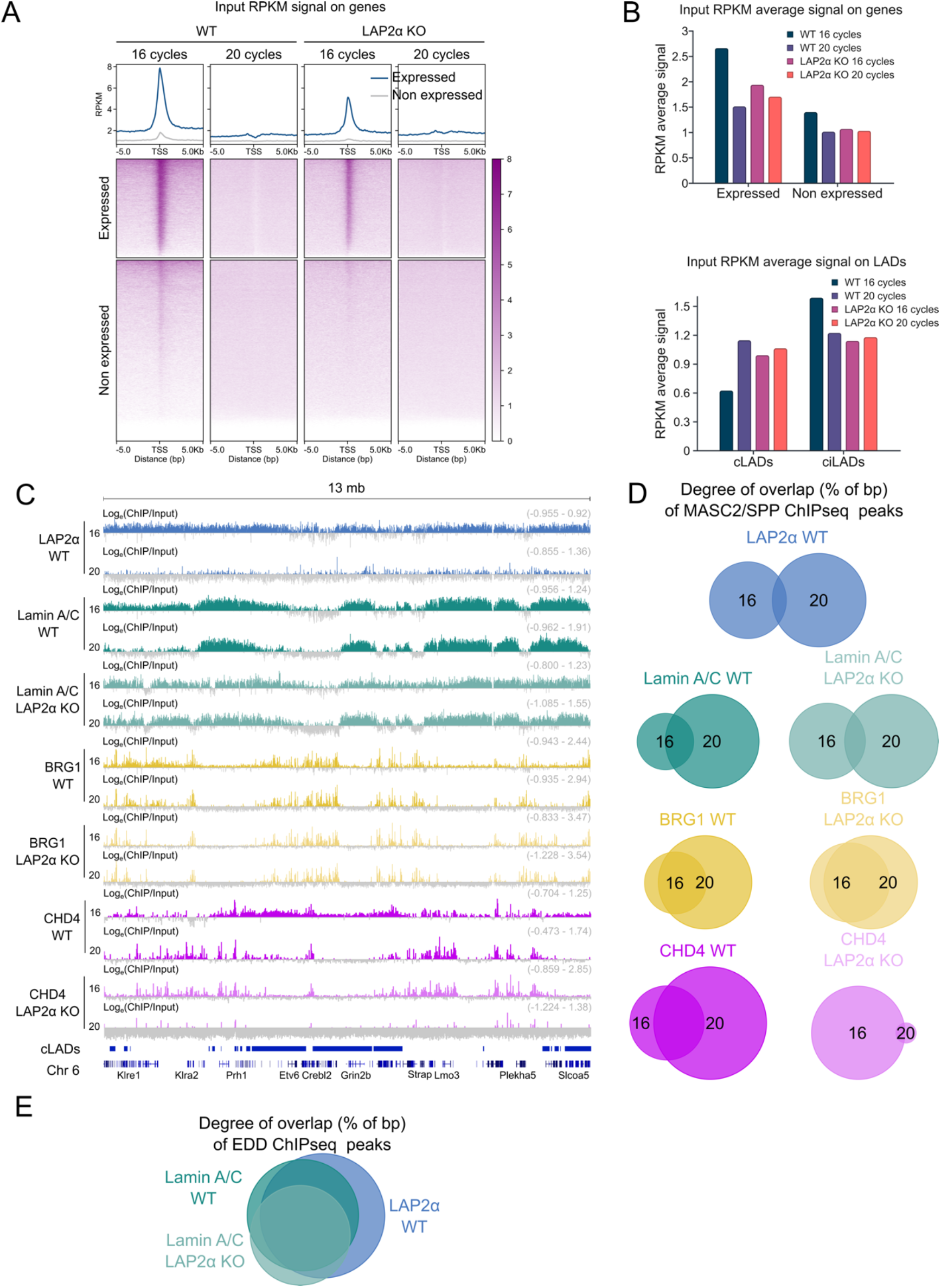
Evaluation of ChIP-seq protocol using 16 and 20 cycles of sonication cycles. **(A)** Heatmaps displaying RPKM signals of input samples from LAP2α wildtype and knockout fibroblasts on expressed genes (blue) and non-expressed genes (gray) +/-5kb from transcription start site (TSS) in 16 and 20 sonication cycles samples. **(B)** Bar graphs presenting average RPKM signals of input samples from LAP2α wildtype and knockout fibroblasts on expressed genes and non-expressed genes in 16 and 20 sonication cycles preparation. **(C)** ChIP-seq analysis was performed in wildtype and LAP2α knockout fibroblasts for LAP2α (blue), lamin A/C (3A6 antibody; turquoise), BRG1 (yellow), and CHD4 (magenta) as indicated. The IGV browser was used to display the log2 ratio of ChIP over input signal from samples sonicated for 16 and 20 cycles, with tracks shown for a 13MB region of mouse chromosome 6. Positive log2 ratio values are shown in color, while negative values are shown in gray. The scale of each log2 ratio track is indicated on the right. Constitutive lamina-associated domains (cLADs) are also annotated. Gene annotations are based on the NCBI reference sequence database. **(D)** Venn diagrams showing the percentage of basepair overlap (% of bp) for ChIP-seq MACS2/SPP peaks of proteins from samples sonicated for 16 and 20 cycles from wildtype and LAP2α knockout fibroblasts, presented in the following order from top to bottom: LAP2α, lamin A/C, BRG1, and CHD4. **(E)** Three-way Venn diagram showing the percentage of base pair overlap (% of bp) for ChIP-seq EDD peaks (16 and 20 samples combined) of LAP2α and lamin A/C in wildtype and LAP2α knockout cells.

**Figure S3.**
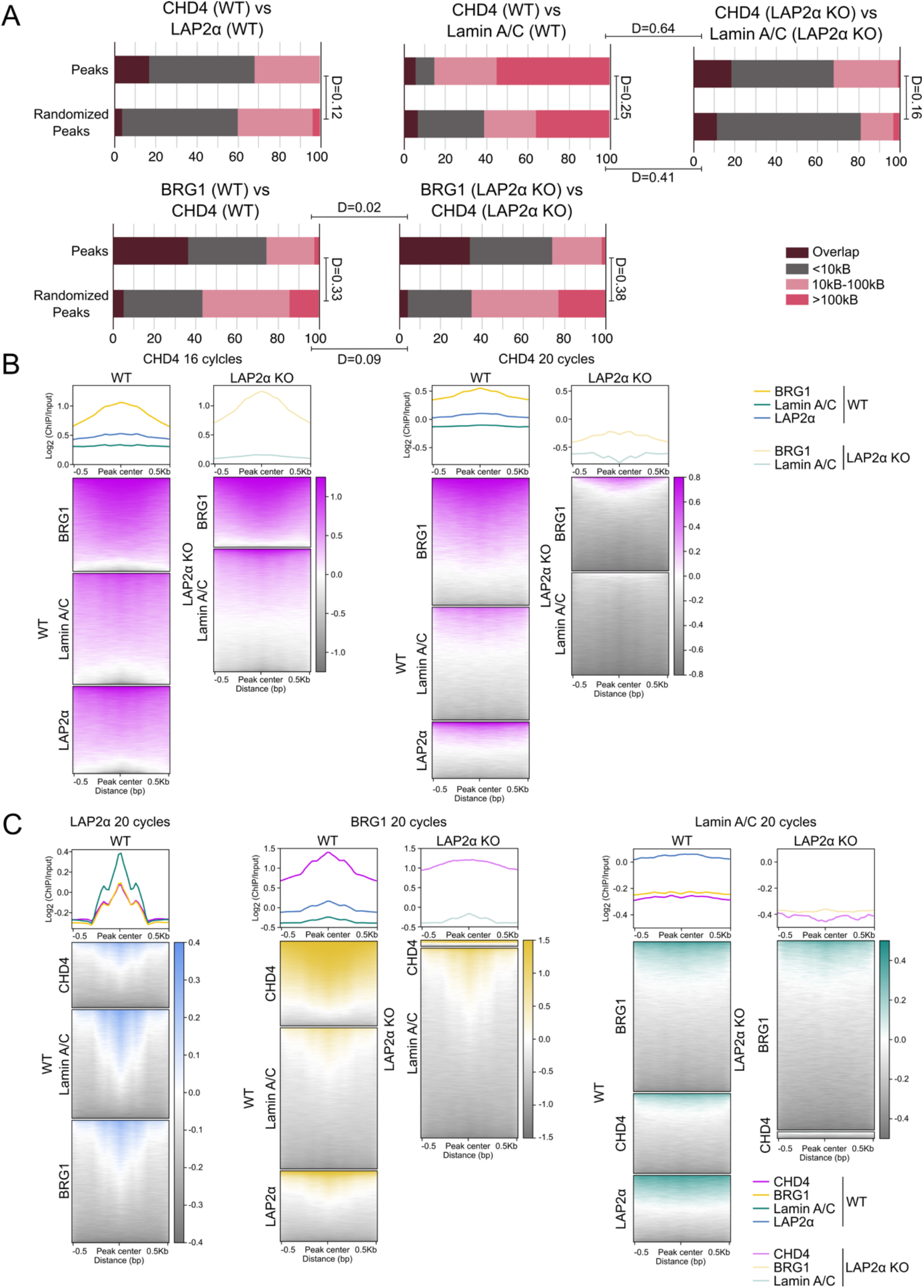
Depletion of LAP2α causes relocation of lamin A/C and chromatin remodelers on chromatin. **(A)** Bar charts showing genomic distances in base pairs between ChIP-seq peaks (16 and 20 sonication cycles peaks combined) for the proteins indicated at the top of each chart in LAP2α wildtype and knockout cells. Distances between peaks of the first protein shown at the top of the chart and randomized peaks for the second protein are included as controls. Differences between distributions are indicated by the D-value from the KS test, representing the maximum distance between the cumulative distributions of the two sets. All KS tests have an adjusted P-value (P adj) below 0.05. **(B, C)** Heatmaps displaying log2 ratio signals (ChIP over input in 16 and 20 sonication samples as indicated) for proteins shown on the top of each heatmap, on combined ChIP-seq peaks +/-0.5kb from peak center for proteins shown on left side of heatmaps in LAP2α wildtype and knockout fibroblasts. Graphs above the heatmaps show the mean log2 ratio signals.

**Figure S4.**
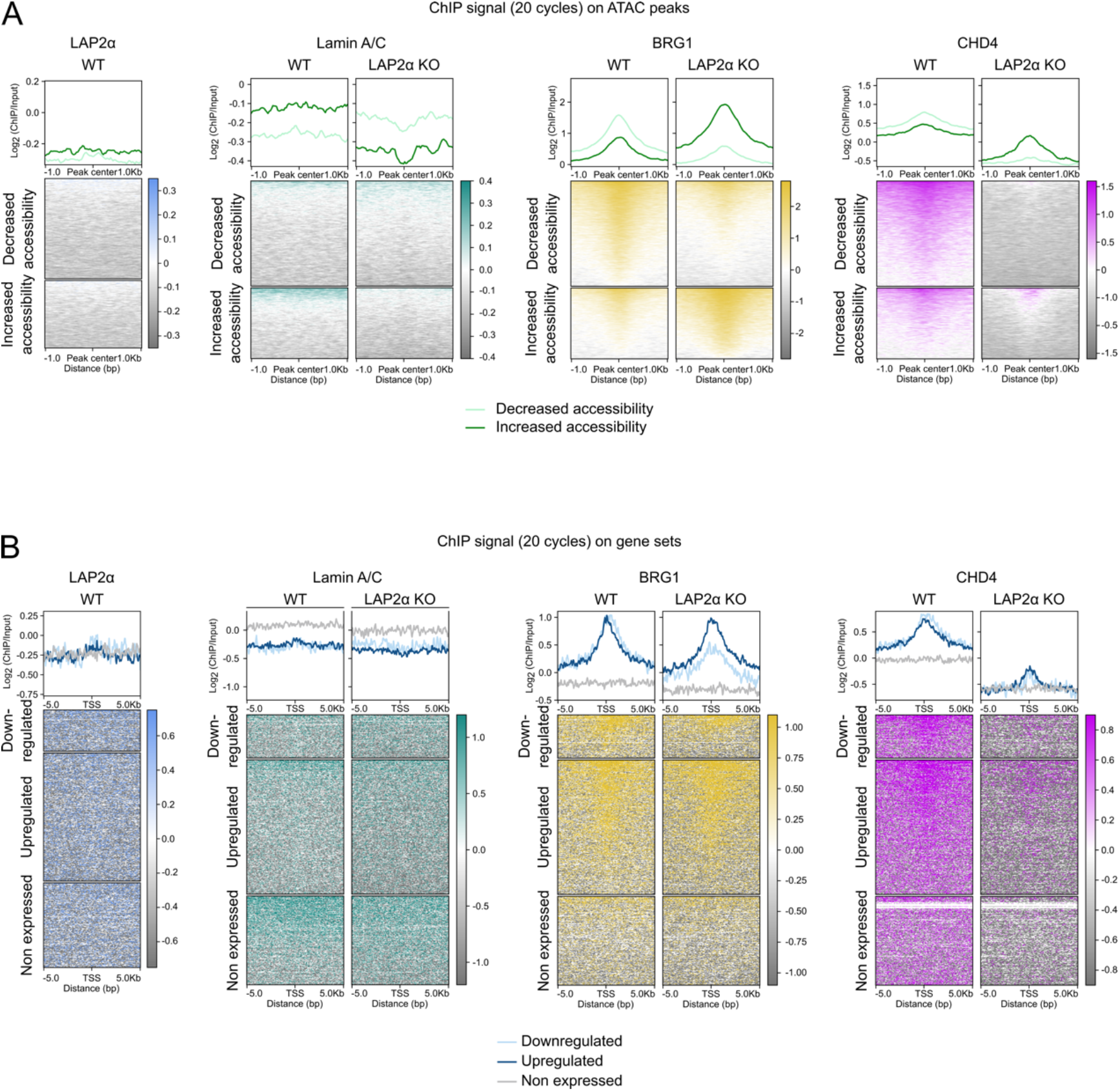
Changes in chromatin accessibility and gene expression correlate with changes in chromatin association of lamin A/C and chromatin remodelers. **(A)** Heatmaps showing log2 ratio signals (ChIP over input) on chromatin regions with decreased accessibility (light green) and increased accessibility (dark green) +/-1.0kb from peak center in LAP2α knockout versus wildtype cells. Log2 ratio signals are presented in the following order from left to right: LAP2α, lamin A/C, BRG1, and CHD4. All four samples sonicated for 20 cycles. Graphs above the heatmaps display the mean log2 ratio signals. **(B)** Heatmaps showing log2 ratio signals (ChIP over input) on downregulated (light blue), upregulated (dark blue) and non-expressed (gray) genes +/-0.5kb from transcription start site (TSS) in LAP2α knockout versus wildtype cells. Log2 ratio signals are presented in the following order from left to right: LAP2α, lamin A/C, BRG1, and CHD4. All four samples sonicated for 20 cycles. Graphs above the heatmaps display the mean log2 ratio signals.

**Figure S5.**
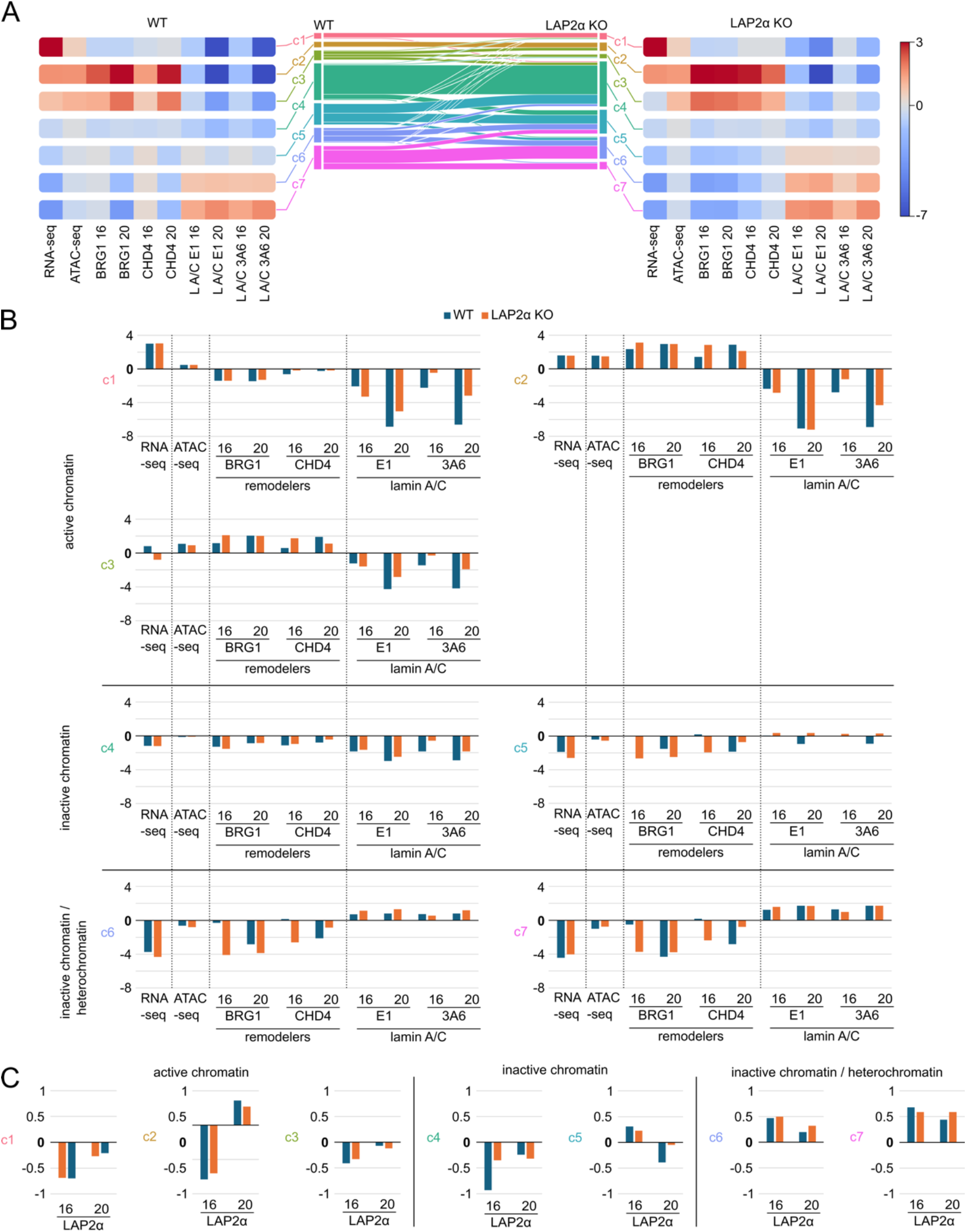
Genome-wide chromatin organization in LAP2α wildtype and KO fibroblasts revealed by clustering. **(A)** 5kb genomic bins covering the entire genome were subdivided into groups of similar signal signatures based on normalized read counts of all datasets listed on the graph using chromHMM. The middle panel displays a plot showing cluster correspondences between wildtype (WT) and LAP2α knockout (KO) cells. The left panel (WT) and right panel (LAP2α KO) present heatmaps showing signal enrichment for individual parameters, including RNA-seq, ATAC-seq, and ChIP-seq (BRG1, CHD4, lamin A/C using E1 and 3A6 antibodies, with 16 and 20 indicating the number of sonication cycles for each sample), relative to the signal in other clusters (log_2_ fold). **(B)** Bar charts displaying signal enrichments from the heatmaps (A) in LAP2α wildtype and knockout cells across all seven clusters. The cluster number is indicated on the left side of each chart. **(C)** Bar charts showing the signal enrichment of LAP2α ChIP-seq signals (with 16 and 20 indicating the number of sonication cycles for each sample) from wildtype cells across all seven clusters in wildtype and LAP2α knockout conditions. The cluster number is indicated on the left side of each chart.

**Figure S6.**
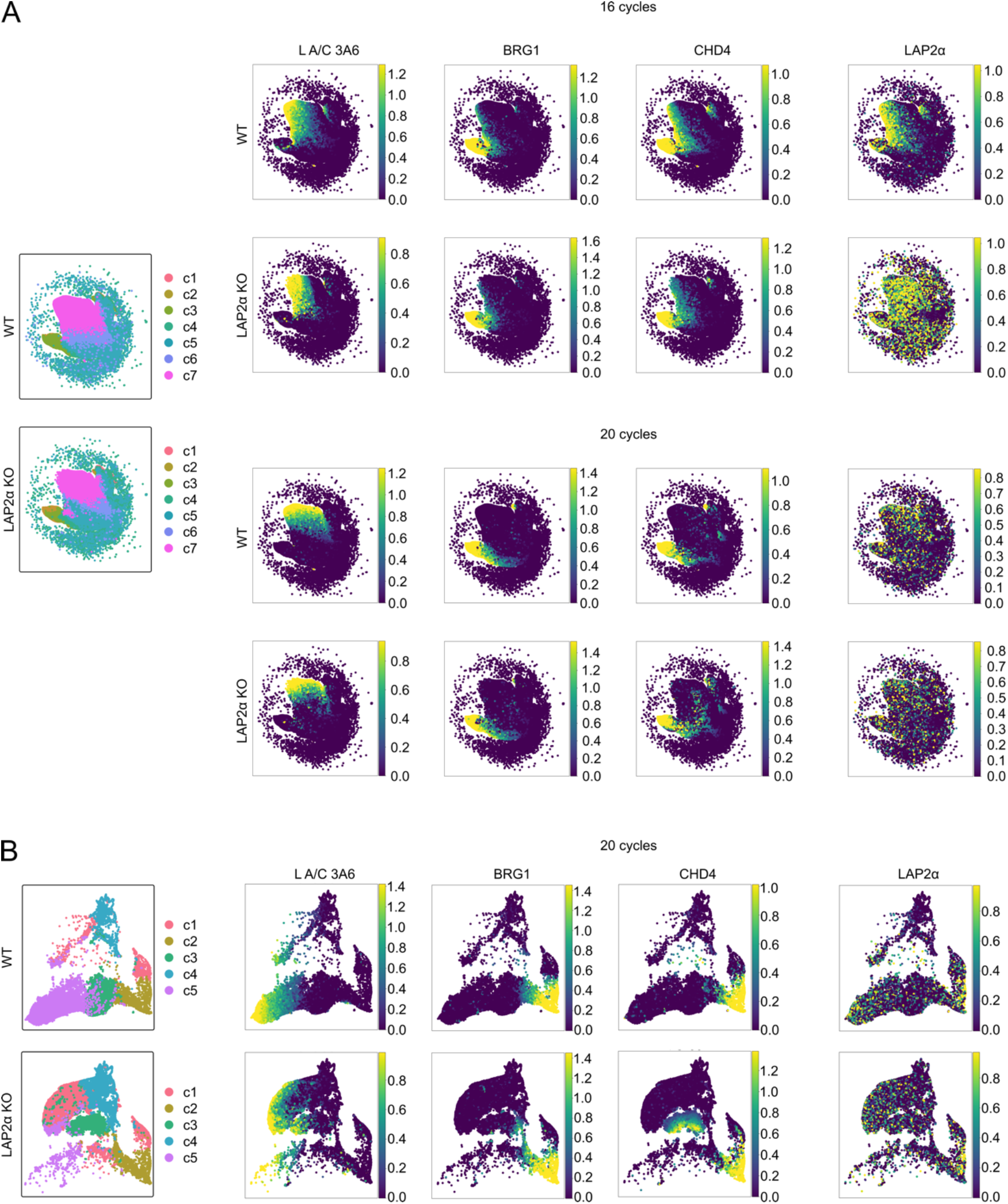
Genome-wide chromatin organization in LAP2α wildtype and KO fibroblasts revealed by Uniform Manifold Approximation and Projection (A, **B)** UMAP graphs showing whole-genome clusters for 16 and 20 cycles sonication) (A) and LAP2α-enriched clusters (20 cycles sonication) (B) for LAP2α wildtype and knockout cells on the left. On the right, heatmaps plotted on the UMAP graphs display signal intensities for lamin A/C, BRG1, CHD4, and LAP2α ChIP-seq signals (max(ChIP – Input RPM, 0)), presented from left to right.

**Additional file 2**

**Supplemental Table S1.**

| Sample | Number of Peaks | Total Peak Length | Average Peak Length | Genome Coverage % |
| --- | --- | --- | --- | --- |
| **MACS2/SPP peaks** |  |  |  |  |
| **16 cycles** |  |  |  |  |
| **LAP2α WT** |  |  |  |  |
| LAP2α | 67600 | 17507210 | 259 | 0.6 |
| BRG1 | 72242 | 43197006 | 598 | 1.6 |
| CHD4 | 49473 | 24062823 | 486 | 0.9 |
| Lamin A/C E1 | 60691 | 14867434 | 245 | 0.5 |
| Lamin A/C 3A6 | 86115 | 23521168 | 273 | 0.9 |
| **LAP2α KO** |  |  |  |  |
| BRG1 | 107358 | 73245978 | 682 | 2.7 |
| CHD4 | 88836 | 52348106 | 589 | 1.9 |
| Lamin A/C E1 | 142774 | 35009536 | 245 | 1.3 |
| Lamin A/C 3A6 | 188210 | 44039359 | 234 | 1.6 |
| **20 cycles** |  |  |  |  |
| **LAP2α WT** |  |  |  |  |
| LAP2α | 91693 | 25704975 | 280 | 0.9 |
| BRG1 | 206126 | 98369486 | 477 | 3.6 |
| CHD4 | 109897 | 54540737 | 496 | 2.0 |
| Lamin A/C E1 | 256803 | 74063282 | 288 | 2.7 |
| Lamin A/C 3A6 | 183166 | 66543988 | 363 | 2.4 |
| **LAP2α KO** |  |  |  |  |
| BRG1 | 235621 | 102081865 | 433 | 3.7 |
| CHD4 | 8387 | 2550218 | 304 | 0.1 |
| Lamin A/C E1 | 390220 | 116213354 | 298 | 4.2 |
| Lamin A/C 3A6 | 257077 | 66012966 | 257 | 2.4 |
| **Merged** |  |  |  |  |
| **LAP2α WT** |  |  |  |  |
| LAP2α | 151219 | 41433796 | 274 | 1.5 |
| BRG1 | 218465 | 107687739 | 493 | 3.9 |
| CHD4 | 129199 | 63359808 | 490 | 2.3 |
| Lamin A/C E1 | 226904 | 79873771 | 352 | 2.9 |
| Lamin A/C 3A6 | 165548 | 55370039 | 334 | 2.0 |
| **LAP2α KO** |  |  |  |  |
| BRG1 | 205784 | 113187462 | 550 | 4.1 |
| CHD4 | 92646 | 53680450 | 579 | 2.0 |
| Lamin A/C E1 | 292093 | 98210303 | 336 | 3.6 |
| Lamin A/C 3A6 | 379003 | 97525031 | 257 | 3.6 |

| EDD peaks |  |  |  |  |
| --- | --- | --- | --- | --- |
| 16 cycles |  |  |  |  |
| **LAP2α WT** |  |  |  |  |
| LAP2α | 312 | 1076178000 | 3449288 | 39.3 |
| Lamin A/C 3A6 | 483 | 967246000 | 2002580 | 35.4 |
| **LAP2α KO** |  |  |  |  |
| Lamin A/C 3A6 | 227 | 351120000 | 1546784 | 12.8 |
| **20 cycles** |  |  |  |  |
| **LAP2α WT** |  |  |  |  |
| LAP2α | 58 | 428214000 | 7383000 | 15.7 |
| Lamin A/C 3A6 | 1084 | 789330000 | 728164 | 28.9 |
| **LAP2α KO** |  |  |  |  |
| Lamin A/C 3A6 | 638 | 792552000 | 1242245 | 29.0 |
| **Merged** |  |  |  |  |
| **LAP2α WT** |  |  |  |  |
| LAP2α | 302 | 1354470000 | 4485000 | 49.5 |
| Lamin A/C 3A6 | 739 | 1078901000 | 1459947 | 39.4 |
| **LAP2α KO** |  |  |  |  |
| Lamin A/C 3A6 | 643 | 870318000 | 1353527 | 31.8 |

