## Additional file 3_uncropped images of Western blots for "Depletion of lamin-associated polypeptide 2alpha leads to chromatin reorganization and binding of A-type lamins to open genomic regions"

### Images of full western blots

A

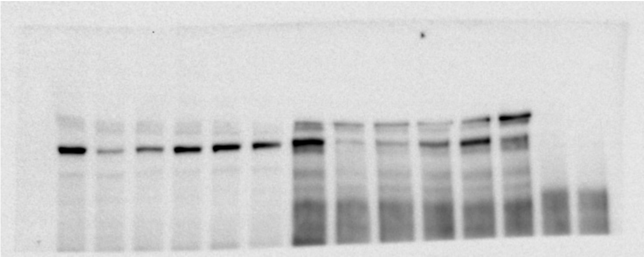

B

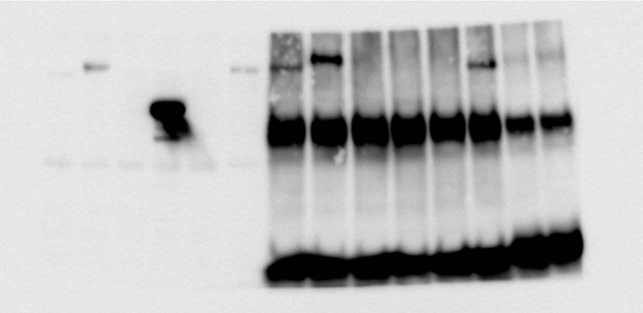

C

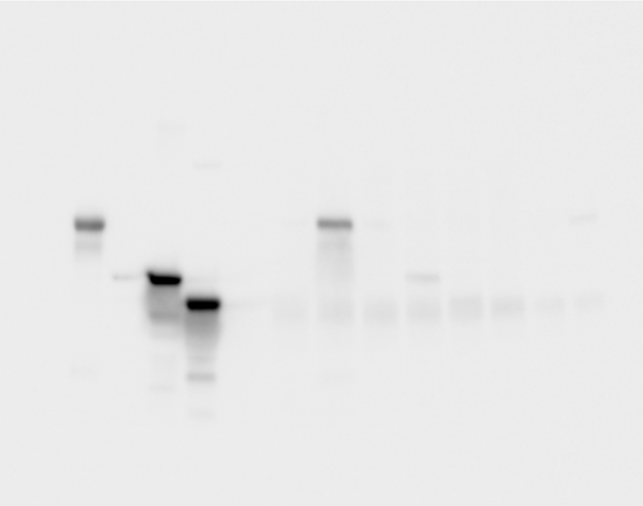

D

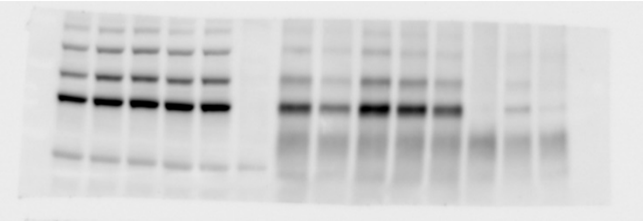

**Western blot for Fig. S1A: (A)** BRG1 **(B)** LAP2α monoclonal **(C)** FLAG-tag **(D)** lamin A/C.

A

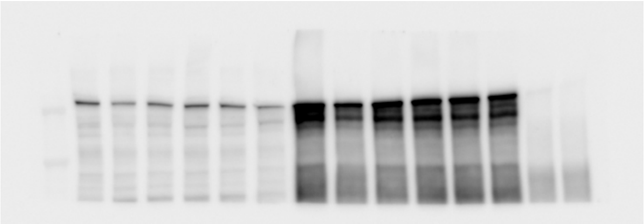

B

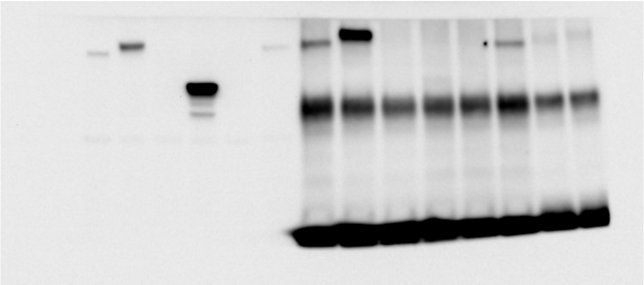

C

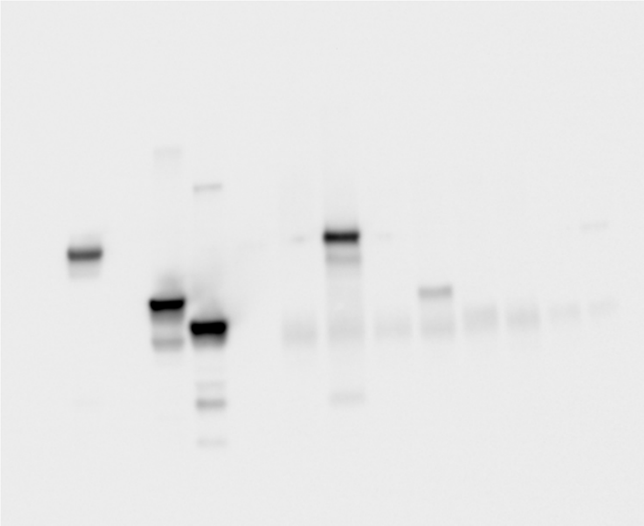

D

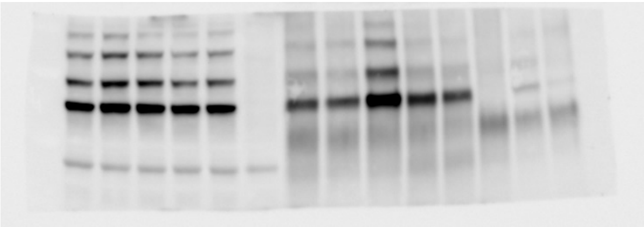

**Western blot for Fig. S1B: (A)** CHD4 **(B)** LAP2α monoclonal **(C)** FLAG-tag **(D)** lamin A/C.
